# Force-induced unfolding versus unbinding of a cellulosomal carbohydrate-binding module from cellulose interface

**DOI:** 10.64898/2026.10.16.759631

**Authors:** Kolattukudy P. Santo, Ryan Murphy, Markus Hackl, Tong Zhong, Shishir P. S. Chundawat, Alexander V. Neimark

**Affiliations:** Department of Chemical and Biochemical Engineering, Rutgers, The State University of New Jersey, 98 Brett Road, Piscataway, New Jersey 08854, United States of America

**Keywords:** acoustic force spectroscopy (AFS), carbohydrate-binding module (CBM), crystalline cellulose, cellulosomal enzyme complex, rupture forces, constant loading rates, protein network analysis, steered molecular dynamics (SMD)

## Abstract

Carbohydrate-binding modules (CBMs) anchor microbial cellulosomes to cellulose surfaces, aiding biomass degradation under mechanical stress. To understand how mechanical pulling affects this process, we combine steered molecular dynamics (SMD) simulations with acoustic force spectroscopy (AFS) to investigate Clostridium thermocellum CBM3a unbinding from cellulose under different pulling configurations (C-versus N-terminus). Our SMD simulations reveal that C-terminal pulling induces substantial CBM3a unfolding prior to unbinding, whereas N-terminal pulling results in detachment with the protein largely intact. Unbinding proceeds via two distinct mechanisms across the planar binding motif: a TRP mode (initiating via TRP135) and a TYR mode (initiating via TYR84). AFS experiments corroborate the SMD findings, showing that C-terminal pulling yields lower rupture forces and a 7.5-fold higher unbinding off-rate compared to N-terminal pulling of CBM3a from cellulose. These results demonstrate that structural asymmetry and pulling orientation critically govern the mechanical stability, unfolding pathways, and cellulose-binding dynamics of cellulosomal proteins.

## 1. Introduction

The stability and function of proteins under mechanical stress is an important biophysical property as many proteins or enzymes are exposed to mechanical stress *in vivo*. For example, kinesin motors move on microtubules in a stepwise fashion, which may induce conformation changes or damage to the microtubule.^1^ In multidomain enzyme complexes, such as microbial cell-surface associated and cell-free secreted cellulosomes degrading cellulosic biomass, mechanical forces acting on one end of the complex can be transmitted throughout the polypeptide chain, and the stability of each domain under external stress is crucial for the function of the complex.^2 3^ Targeting of enzymes to specific carbohydrate-ligands is often driven by carbohydrate-binding modules (CBMs) that facilitates the hydrolysis of associated polysaccharides by one or more tethered catalytic domains (CD).^3, 4^ Depending on the sequence of the cellulosomal complex (e.g., cipA from C. thermocellumn), the CBM domain associated with cellulose can experience asymmetric pulling forces via either its N-terminus versus C-terminus due to tethering of the cellulose-bound complex also to the cell surface that can experience shear forces via microbial cell motility or other microenvironment-associated motions. However, the subtle interplay of interfacial CBM binding-unbinding to insoluble polysaccharides as a function of applied shear force and its impact on CD activity is still poorly understood. Therefore, in recent years, advanced single-molecule force spectroscopy techniques (e.g., Atomic Force Microscopy, Acoustic and Optical Tweezers Force Spectroscopy) have been applied to shed light on the unbinding behavior of both cellulosomal^5 6–9^and non-cellulosomal^10–12^ enzyme systems under force. However, in some experimental designs, the dissociation of a protein complex from its ligand may be induced by partial unfolding of protein domain.^10^ The pulling configuration-pulling from the C-versus N-terminus-may impact the obtained rupture forces^11^, as well as the degree of unfolding,^12^ thus careful attention must be paid in the design of rupture force experiments and interpretation of results within relevant context of the biological problem.

It is well known that the presence of the planar aromatic residues on the CBM-cellulose binding interface plays a pivotal role in the strength of the interaction.^12–18^ Since cell-surface associated cellulosomal complexes often degrade insoluble cellulosic substrates in native environments that experience high interfacial shear forces, it’s not unsurprising that these protein complexes have evolved to withstand unbinding at high forces and/or are sensitive to only unbinding in specific modes upon force application. Previous experimental work on the cellulosomal scaffold protein (cipA), cohesion domain, that enables engagement with the dockerin domain containing CDs are relatively stable interactions that only unfold at ruptures forces exceeding 140 pN.^19, 20^ However, similar studies to measure the unbinding rupture forces and modes for CBM-cellulose have only recently been explored. Our recent acoustic force spectroscopy (AFS) study had demonstrated how there were clear differences in the single-molecule rupture force distribution and bond lifetimes that revealed distinct differences between CBM3a native protein and its Y84A (NOTE: Y84 was reported as Y67 in that previous study) mutant.^9^ Interestingly, in that previous work, the multimodal shape of rupture force distributions observed suggested that that CBM3a binding motif has preferred binding orientations on the cellulose surface. Previous MD simulations conducted also showed that in the absence of Y84 the whole CBM can tilt towards W135 position, resulting in more frequent H-bond formation between polar residues and cellulose. Therefore, the multimodal rupture force distribution observed for the native CBM3a unbinding from cellulose was hypothesized to arise largely due to distinct CBM-cellulose binding orientations and unbinding modes possible. However, the molecular mechanisms of CBM unfolding and its impact on dissociation from cellulose surface are still not well understood.

Molecular dynamic (MD) simulations have illustrated the critical impact of aromatic residues on CBM-Cellulose interactions,^8, 18^ highlighting the need for further exploration into the molecular basis for CBM rupture force trends. MD simulations have been used for estimating free energies of CBM binding to carbohydrate polymer surfaces. Binding free energies of Cel7A and Cel6A CBMs as well as their mutants, have been found to differ between parallel and antiparallel orientations^21, 22^ on various cellulose surfaces, while CBM20^23^ has shown a greater affinity when binding to the helical structures of starch. The steered molecular dynamics (SMD) approach^24^ has been used to examine the response of macromolecules to mechanical stress, as well as the stability of protein-ligand and protein-protein complexes, aiding in rational drug design.^25–28^ In these simulations, the protein is pulled by a constant or harmonic (umbrella) force at a constant velocity, which can elucidate protein unfolding behavior under an external pulling force.^29–33^ However, such studies have not been performed to examine the unbinding behavior of CBMs from carbohydrate surfaces like cellulose under different pulling configurations.

In this study, by using advanced force spectroscopy SMD simulations and AFS experiments, we aim to identify the modes by which a well-known cellulosomal CBM3a from *C1ostridium thermocellum*,^4,7,8,19–22^ unbinds from the cellulose surface, and the impact of pulling the protein from its N-versus C-terminus as well as the applied force loading rate on the unbinding behavior. To elucidate the nature of the CBM-cellulose bond stability, we established a protocol to conduct single-molecule rupture force experiments *in silico*, in which the protein is pulled at a constant loading rate. We demonstrate that the pulling configuration strongly influences the protein conformations and rupture forces at the various studied loading rates. This work highlights the importance of combining *in-silico and in-vitro* experiments to elucidate force-induced dissociation mechanism of protein-carbohydrate complexes.

## 2. Methods

### 2.1 Molecular Dynamics Simulations

#### CBM3a Adsorbed on a Cellulose Fibril

CBM3a protein structure used in this work corresponds to the PDBID:4JO5, that has residues numbered from 12 to 181 (170 residues in total). The protein-cellulose model system is constructed as described earlier (see Supplementary Information of our previously published work^9^). The cellulose fibril (called CelF from hereon) is modeled as a bundle of 41 chains, each 12 monomers long. The protein is placed on the cellulose surface, with residues Y84, H74, R129 and W135 positioned close to the surface of the cellulose fiber **(Figure 1a)**. The *Glycam06*^*34*^ force field is used for modeling the cellulose chains while *amber99sb-ildn*^*35*^ *force* field is used for CBM3a. The forcefield parameters for the cellulose fiber are obtained using *Antechamber module* of the *AmberTools*, and are converted to GROMACS format using the *Acpype*^*36*^ Python utility.

**Figure 1.**
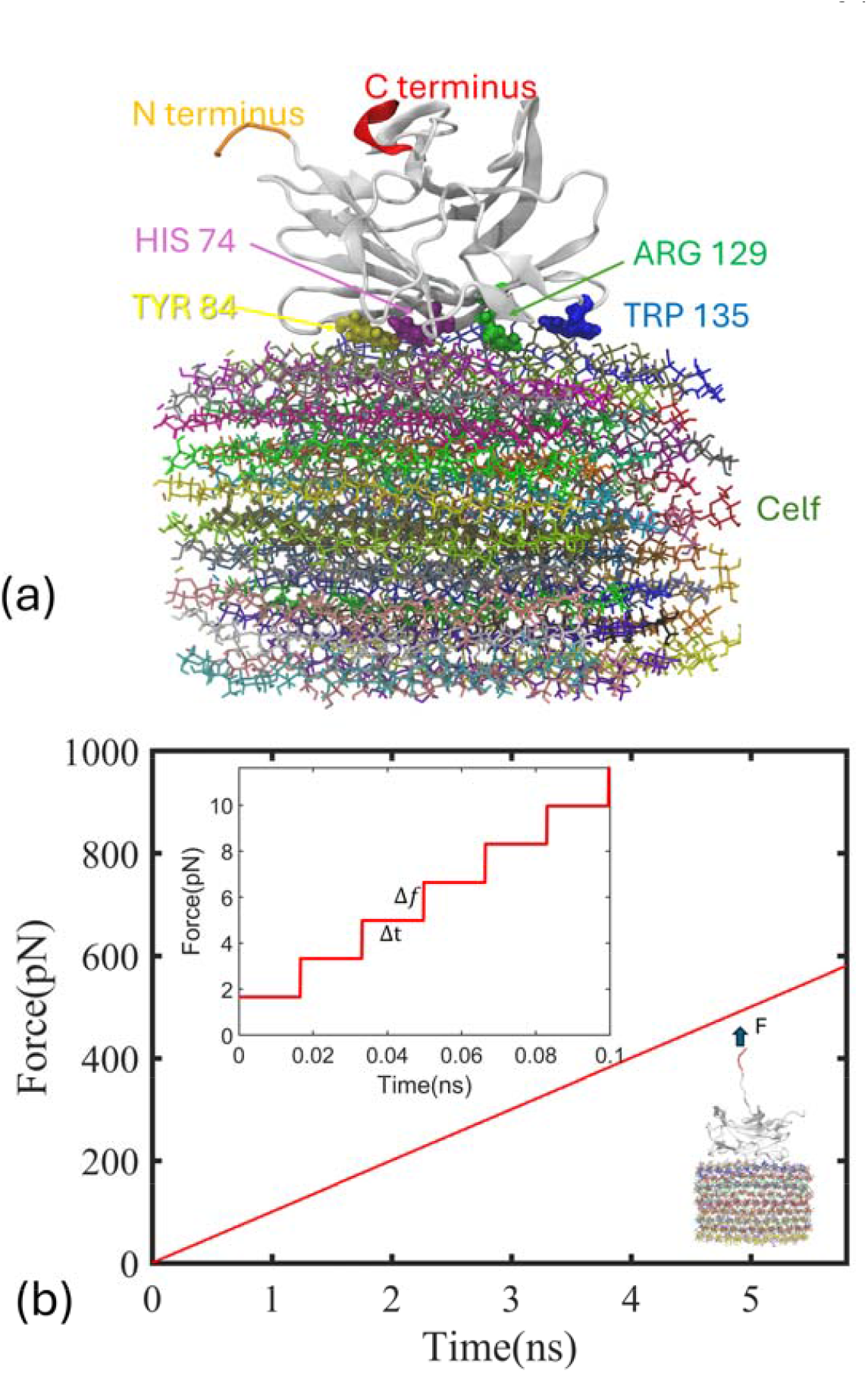
Overview of CBM3a-Cellulose MD Methodology: (a) The equilibrated CBM3a protein structure on the cellulose fiber (CelF) surface. Each polymer in the cellulose fibril is colored differently. (b) The variation of force in the constant loading rate pulling simulations reported in this work. The force is increased in steps of at every equal sized interval of Δ*t* (inset). This leads to a linear increase in force with time that tends to be continuous as Δ*f*, Δ → 0.

The simulations are performed with GROMACS 2021.6.^37^ The system size is chosen to be long, ~50 nm, in the z direction, in order to allow extension of the protein in this direction of pulling. The protein-CelF system is solvated with TIP3P water.^38^ Potassium (K+) and chloride (Cl) ions are added to neutralize the protein charge and to adjust the ionic strength of the system to 0.15 M. The system is energy-minimized using the steepest decent algorithm and then relaxed with a 200 ps NVT simulation with position restraints applied to both the protein and CelF. This is followed by another 100 ps NVT simulation with position restraints only on C1, C3, and C4 atoms of the terminal glucose units of the cellulose chains. Water hydrogens are constrained using with SETTLE,^39^ while other covalent bonds involving hydrogens are constrained with LINCS^40^ algorithm. The non-bonded interactions are cut off at 1.2 nm, and long-range electrostatic interactions are computed using the smooth particle mesh Ewald^41, 42^ (SPME) algorithm. The temperature is controlled at 31 OK using the v-rescale^43^ thermostat, and a time step of 2 fs is used. The system is further relaxed by running NPT simulations: a 200 ps simulation with position restraints on both the protein and CelF, followed by a 1 ns simulation with only the terminal glucose carbon atoms of the cellulose chains restrained. The pressure is controlled with Berendsen barostat at 1 bar with coupling constant 2 ps and compressibility 4.5 × 10^−5^ bar^−1^. The production run is conducted for 10 ns at NPT conditions with position restraints applied only on the C1, C3 and C4 carbon atoms of the terminal glucose units. The temperature is controlled at 310K using the Nose-Hoover^44, 45^ with coupling constant 2 ps. To control pressure at 1 bar, the Parrinello-Rahman barostat^46, 47^ is employed with coupling constant 4 ps. The final system size is 8.03 × 6.03 × 50.2 *nm*^*3*^ with 237863 atoms.

#### Constant Loading Rate Steered MD Pulling Simulations

We pull the protein from either N-or C-terminus end at a constant loading rate, mimicking the experimental procedure. To perform the pulling, we use the GROMACS *pullcode* utility, which can execute constant force/velocity pulling on the pull group from the center of mass from a group of atoms that are restrained at respective positions with harmonic bonds. Keeping the heavy atoms in the CelF restrained in space, we perform pulling from both C-terminal and N-terminal ends of the protein. The pull groups are chosen to be five consecutive residues at each terminus; the C-terminus is pulled by applying force on residues 177-181, while the N-terminus is pulled with the residues 12-16. To implement pulling at constant loading rate in which the force increases linearly with time, we devised the following strategy: A constant force is applied on the pull group along the pulling coordinate for a small-time interval Δ*t*, and then the force is increased in small increments Δ*f* during the successive time intervals. This results in a force being applied on the pull groups that increases linearly with time, the loading rate is given by,

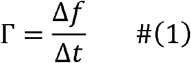

Γ is constant as long as both Δ*f* and Δ*t* remains the same for all intervals in the simulation. We performed pulling simulations keeping Δ*t* = 16.6 ps same for all simulations but varying Δ*f* to apply loading rates within the range of Γ = 0.1 – 0.5 N/s. The simulations are stopped atthe end of every Δ*t* interval and then restarted, continuing from the last state with force increased by Δ*f*. Several hundreds of such steps are involved in a simulation until the CBM3a gets unbound from the cellulose. As Δ*t* is chosen to be much smaller compared to the total length of the simulations (~ ns), the force vs time curve appears linear on the ns timescale (Figure 1b) effectively demonstrating constant loading rate pulling. Note that Δ*t* = 16.6 ps is chosen only for convenience and can be varied, however, it has to be chosen sufficiently small, as making it large will affect the continuity of the constant loading.

#### Protein Network Analysis

Network analysis is a powerful tool to elucidate strongly linked residue communities and their intra- and inter-connectivity.^48, 49^ We performed network analysis of the CBM3a equilibrated trajectory (last 5 ns) using the recently developed Dynamic Network Analysis (DNA) approach.^49^ In this analysis, the protein structure is reduced to ‘nodes’ connected with ‘edges’, where each node represents a residue at the *α*-carbon. The edges between the nodes represent a residue-residue contact determined by a cutoff for the minimum distance between the heavy atoms of the residues, taken here to be 4.5 Å. Additionally, each edge is assigned a weight given by the generalized correlation coefficient, which represents the correlation between the positional variations of a pair of nodes over the trajectory window. The communities are identified through modularity optimization, which maximizes the density of edges within communities compared to the edges between communities.^50^

### 2.2 Characterization of CBM3a-Cellulose Binding by Acoustic Force Spectroscopy

#### C1oning

DNA fragment gBlocks and relevant primers for cloning CBM3a linked to GFP (Green Fluorescent Protein) with Avi- and His-tags at the C and N-terminus, respectively, were purchased from Integrated DNA Technologies (Coralville, Iowa). Gene fragments are cloned into a pEC vector by Gibson assembly using a New England Biolabs Builder kit (Ipswich, MA). Plasmids are transfected into BL21 DE3 cells with the BirA plasmid for in-vivo biotinylation (as described in our published work^51^). Colonies are grown on LB-agar plates with kanamycin for selection. Colony screening was performed to confirm inserted gene of interest and select colonies were grown in 10 mL culture tubes for plasmid DNA extraction and sequencing. Plasmid extraction was performed using IBI Scientific Plasmid mini-prep kits (Dubuque, Iowa). Sequences were confirmed with Sanger sequencing by Azenta-Genewiz (Piscataway, NJ). All relevant sequence and primer information is provided in the SI document.

#### Expression and Purification ofCBM3a

The details of the CBM3a production methodology are described in a previous paper.^51^ Briefly, colonies with pEC-CBM3a-GFP-Avi-His (C-terminus pulling construct) and pEC-Avi-His-GFP-CBM3a (N-terminus pulling construct) were grown in 10 mL LB media with carbenicillin overnight at 37°C with a shaking speed of 200 rpm. Next, 5 mL of starter culture volume is transferred to 50 mL LB media with carbenicillin and incubated at the same conditions to an OD600 of ~0.3. At OD 0.3, protein expression is induced by addition of IPTG at a final concentration of 1 mM. Biotin is also added to a final concentration of 0.2 mM. Biotin is attached *in vivo* to the GFP-CBM, at the Avi tag site located at either terminus of the protein by BirA enzyme over-expressed in *E. coli*.

Harvested cell pellets were resuspended in 7 mL of cell lysis buffer (10 mM PB at pH 7.4 containing 100 mM 149 sodium chloride, 20% v/v glycerol and 0.3% v/v Triton X-100) with Lysozyme and a Protease inhibitor cocktail of E64 protease inhibitor in 0.5 mM Benzamidine, 1 mM EDTA. Cells were lysed by sonication 10 seconds on, 30 seconds off, for 5 minutes total. Lysed cells were centrifuged for 40 minutes at4°C and 3000 rpm and the supernatant was decanted for further processing.

GFP tagged CBM3as were purified by IMAC using Ni-NTA magnetic beads. C1arified lysates were incubated with 1 mL of Ni-NTA magnetic beads at4°C for 60 minutes with continuous gentle shaking. The supernatant is removed, and the magnetic beads are incubated with 20 mL lysis buffer for 20 min at4°C with gentle shaking. The supernatant is again removed, and the beads are washed the same way with 10 mM Sodium Phosphate, pH 7.4. The beads are then washed the same way, twice with IMAC-A buffer (100 mM MOPS, 10 mM imidazole, 500 mM sodium chloride and 0.1 % v/v Triton X-100 at pH 7.4), and twice with 5% IMAC-B (100 mM MOPS, 500 mM imidazole and 500 mM sodium chloride at pH 7.4) 95% IMAC-A buffer. Finally, the GFP tagged CBMs were eluted in 1 mL of 100% IMAC-B for 15 min at4°C. Proteins are buffer exchanged into 100 mM MOPs, 500 mM sodium chloride using 30kDa cutoff, regenerated cellulose filter columns and centrifuging at 3000 rpm for long term storage at −80°C.

#### AFSexperiments

All experiments were performed on LUMICKS AFS system and z-Movi microfluidic chips (Amsterdam, The Netherlands) functionalized with nanocrystalline cellulose (NCC) derived by treatment with H_2_SO_4_ as described in our previous study.^51^ Briefly, z-Movi chip surface passivation was performed with 10 mM Sodium Phosphate, pH 7.4, buffer supplemented with 2.5 mg/mL of BSA and Casein proteins, and 5.6 mg/mL of Pluronic surfactant. Calibration of the acoustic forces experienced by each polystyrene bead inside the AFS chip was done using a custom in-house MATLAB based analysis protocol and graphical user interface, as was previously described.^51^ All rupture force at various loading rates were performed with 10 mM Sodium Phosphate, pH 7.4 as working buffer.

Biotinylated GFP tagged CBMs (GFP-CBMs) were incubated with streptavidin coated 5 µm polystyrene (PS) beads from Spherotech (Lake Forest, IL), at a concentration of 1.5 µM GFP-CBMs, in 0.17% w/v of PS beads. The protein concentration was chosen based on previous work where rupture forces remain unchanged at higher concentrations.^51^ The binding interactions of PS immobilized CBM3a was evaluated using our effector bead AFS assay methodology as described by Hackl et al.^51^ Biotinylated GFP was linked to CBM3a at either its C- or N-terminus, which was then functionalized onto streptavidin-coated 5 µm polystyrene beads via a biotin-streptavidin linkage (see **Figure 2** for overview of assay setup and workflow). Image acquisition was done at 40 Hz using a lOx Nikon Plan Fluor objective, with a 0.30 numerical aperture. This configuration allows the application of controlled forces on CBM3a via the tethered PS beads and ensures precise tracking of CBM functionalized PS beads and the determination of rupture forces for bead unbinding from cellulose surface. To determine binding parameters of CBMs to nanocrystalline cellulose, rupture force experiments are conducted in duplicate at four force loading rates, across four orders of magnitude (0.1 pN/s −100 pN/s). Beads were allowed to bind on to the NCC surface for 15 minutes, prior to a 20-minute washing step, with 10 mM Sodium phosphate, pH 7.4, buffer at a flow rate of 3 µL /min, with an applied force of ~1 pN. The Bell-Evans (BE) model^52^ for analyzing the rupture force dataset was applied to extrapolate apparent k’_Off_ and transition state distance Δ*x*′ values.

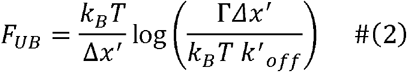

**Figure 2.**
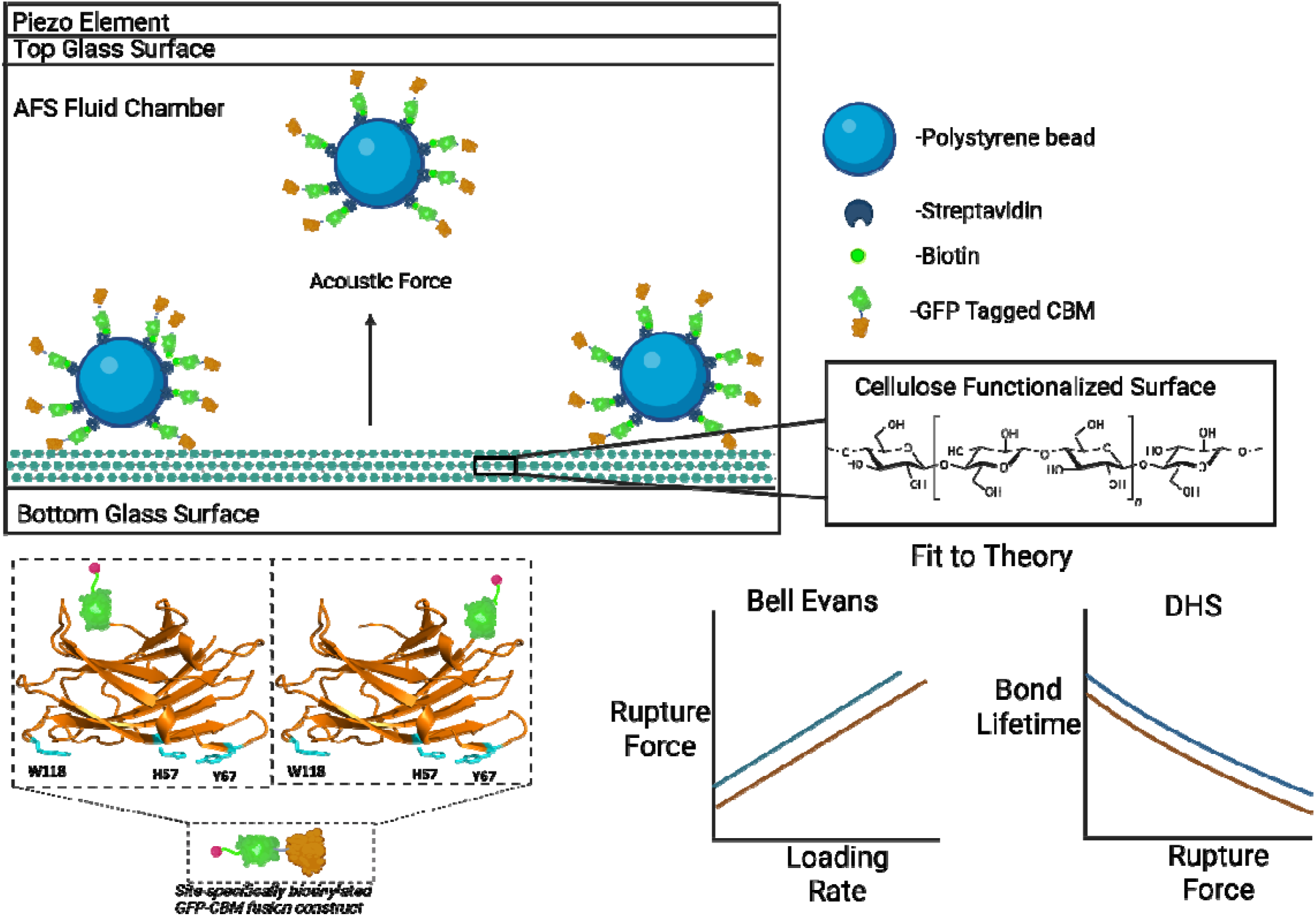
Overview of CBM3a-CelluloseAFS Binding Assay Methodology: CBM3a (orange color) linked to GFP (green color) at either the C- or the N-terminus with biotin tag (filled cirC1e symbol), is functionalized directly onto polystyrene beads with streptavidin via biotin-streptavidin interactions to mimic *C. thermocellum* cellulolytic bacteria with cell surface displayed cellulosomal proteins. The CBM3a displayed proteins on beads were bound and then pulled vertically away from a surface of nanocrystalline cellulose fibers immobilized inside an Acoustic Force Spectroscopy (AFS) z-Movi microfluidic chip using the LUMICKS AFS system. The resulting measured rupture forces and bond lifetimes at various AFS loading rates were fit to various single-molecule force spectroscopy theory (e.g., Bell Evans and Dudko-Hummer-Sudko or DHS models) to determine CBM3a-cellulose bond properties. Here, CBM3a binding motif with key aromatic residues (Y67, H57, and W118) are numbered based on prior work and are equivalent to Y84, H74, and W135 numbering scheme used in this study, respectively.

Here, *k*_*B*_ is the Boltzmann’s constant and *T* is the temperature. All experiments were performed at room temperature (approximately 300K).

## 3. Results and Discussion

### 3.1 Steered MD CBM3a Pulling Simulations from Cellulose Fibril Surface

#### Force vs extension curves

We performed SMD simulations of the CBM3a protein being pulled from its two different termini at a constant loading rate Γ, ranging from 0.1 to 0.5 N/s. As the protein is pulled in the z-direction away from the CelF surface, the force applied increases as a function of time, F(t) = F_o_ + Γt, with initial force chosen to be *F*_*o*_ = 0, and the pulled protein moves in the *z-* direction causing an extension *z*_*ex*_*(t) = z(t) −* z(0) in the polypeptide chain, where *z(t)* is the position of the center of mass of the terminal pull-group residues (see Methods) at time *t*. At some point, the protein unbinds from the CelF ligand surface, characterized by an unbinding force F_UB_(Γ) and a corresponding extension *z*_*UB*_ (Γ), both of which are functions of the loading rate. From simulations, we determined both *F*_*UB*_ and *z*_*UB*_ by monitoring the number of CelF-CBM3a atomic pair contacts over time, with unbinding determined as the point when such pair contacts reduce to zero. See **Supplementary Information Figure S1** for representative SMD datasets.

Figure 3. presents key observations from SMD simulations of the CBM3a unbinding process during either terminal pulling. As shown in the representative snapshots of the unbinding process **(Figure 3a-b)**, pulling the CBM3a from its C-terminus leads to considerable unwinding or unfolding of the protein at all loading rates analyzed. When pulled from N-terminus at lower loading rates, however, CBM3a detaches from CelF surface without any unfolding, although partial unwinding occurs at higher loading rates. **Figures 3c-d** illustrate the effects of pulling from different ends on the CBM3a extension, as a function of force at all loading rates considered. **Figure 3c** shows that in the C-terminal pulling mode, CBM3a easily begins to unwind at lower forces and lower loading rates, before completely detaching from the cellulose surface. In contrast, **Figure 3d** shows that in the N-terminal pulling mode, CBM3a remains compact at low loading rates and does not unwind as the force increases until it eventually detaches. When pulled from the N-terminus, the protein unbinds without unfolding at loading rates of 0.1 N/s, 0.2 N/s, and 0.3 N/s, while partial unwinding is observed at higher loading rates, specifically at 0.4 N/s and 0.5 N/s. However, even at higher loading rates, unfolding does not occur in the N-terminal pulling mode until a large force is reached **(Figure 3d)**.The unwinding occurs at Γ = 0.4 *N/s*, only when the force exceeds the unbinding force *F*_*UB*_ observed at loading rate 0.3 N/s. This indicates that, in the N-terminal pulling mode, the protein tends to remain intact at small, and especially, at experimentally relevant, lower loading rates.

**Figure 3.**
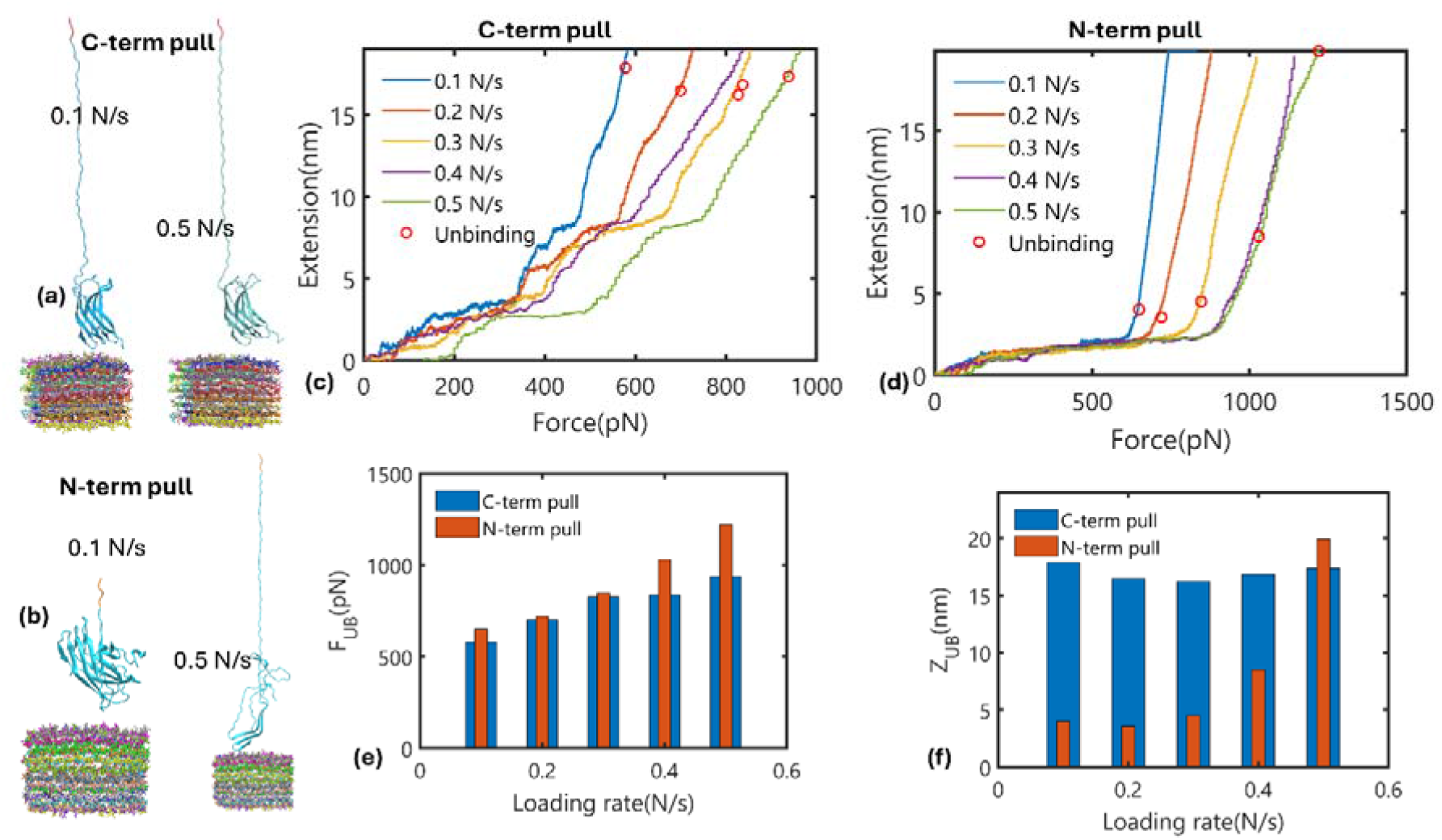
SMD simulation of CBM3a unfolding/unbinding from cellulose surface: Snapshots of the CBM3a unbinding from cellulose surface after being pulled from the (a) C-terminus and (b) N-terminus, at different representative loading rates is shown here. Each polymer in the cellulose fibril is colored differently. The pulled protein residues are colored in red (C-terminus) and orange (N-terminus), while the rest of the protein in colored in cyan. The force versus extension curves at different loading rates for different pulling configurations (c-d) and (e) unbinding forces and corresponding extensions (f) at different pulling configurations are shown here.

**Figure 3e-f** illustrates the unbinding forces *F*_*UB*_ and the corresponding extension *z*_*UB*_ as a function of the loading rate. The dependence of *F*_*UB*_ on the loading rate varies across different pulling modes. In the C-terminal pulling mode, the stepwise behavior in **Figure 3c** indicates the existence of multiple energy barriers for unwinding to overcome, before unbinding. In contrast, the N-terminal pulling mode involves only a single step of unbinding, and the unbinding forces increase with the loading rate as expected in a barrier crossing event. Although some unwinding occurs at very large loading rates, it happens at much high forces unlike in the C-terminal pulling simulations, and therefore this is unlikely to be rate-limiting under relevant experimental conditions. In the C-terminal pulling, *z*_*UB*_ is large ~17 *nm* and it remains practically independent of the loading rate **(Figure 3e)**, indicating greater unwinding flexibility of the protein in this pulling mode. In the N-terminal pulling mode, the low *z*_*UB*_ value (~4 *nm)* at smaller forces indicates that the protein remains rigid, leading to unbinding without significant unwinding in this mode.

#### Unbinding mechanisms

The unbinding of CBM3a protein from the cellulose surface may involve several mechanisms, especially when pulled from a specific terminus. In our recent single-molecule force spectroscopy experimental study on CBM3a unbinding from cellulose surface,^9^ the pulling from the N-termini of the protein was assisted by thermal activation over macroscopic time scales, and hence the unbinding occurs at much lower rupture forces than what is measured in MD simulations. CBM3a is bound to the cellulose surface, with key contact residues on the binding motif (namely TYR 84, HIS 74, ARG129 and TRP 135) in close contact with the cellulose surface (Figure 1a). Since TYR 84 and TRP 135 are at either side edges of the CBM3a planar binding motif, unbinding of any of these residues becomes rate-limiting and decisive of the protein unbinding mode, as also reported in our recent work.^53^ Both residues were found to be in close contact with cellulose atoms; the minimum distances, *d*_*min*_, between a cellulose atom and these residues, averaged over the last 5 ns of the equilibration simulation are found to be 0.23 nm for both. The average numbers of atom pair contacts with CelF atoms that are less than 0.35 nm are found to be 16 and 14, respectively, for TRP 135 and TYR 84. The closest pair contact of TYR84 remains with one cellulose chain, (chain ID:36) over 98% of time during the equilibration (3% of the time with chain ID 19), while TRP135 remains in closest contact with two chains over 5 ns of time; more closely with chain ID:19 (61% of the time) and with chain ID:36 (39% of time). Our recent force spectroscopy experimental study also showed that mutation at TRP 135 position causes a significant reduction in the bond lifetime unlike mutation made at the TYR 84 position when CBM3a is pulled from its N-termini.^53^

As shown in **Figure 4a**, the unbinding of CBM3a from the cellulose surface occurs with the residue TRP 135 unbinding first and TYR 84 last, when pulled from the C-terminus, at all loading rates simulated. Considerable unwinding of the protein occurs prior to TRP unbinding. **Figure 4b** depicts the order of the residue unbinding in terms of the *d*_*min*_ as a function of the simulation time, showing that TRP 135 (solid lines) unbinds prior to TYR 84 (dashed lines) at all loading rates. The situation is quite different in the case of N-terminal pulling; there are two modes of unbinding depending on whether **TRP** 135 or TYR 84 unbinds first **(Figure 4c)**. Interestingly, the *TYR mode of* unbinding occurs with the protein structure being intact, without any unfolding occurring, while *TRP mode occurs* with considerable unwinding/unfolding of the protein structure always. Note that with C-terminal pulling only *TRP mode* of unbinding is found to occur, while with N-terminal pulling, *TRP mode* occurs only at higher loading rates. **Figure 4d** describes the minimum distances of the residues with CelF atoms as a function of time, showing order of the residue unbinding.

**Figure 4.**
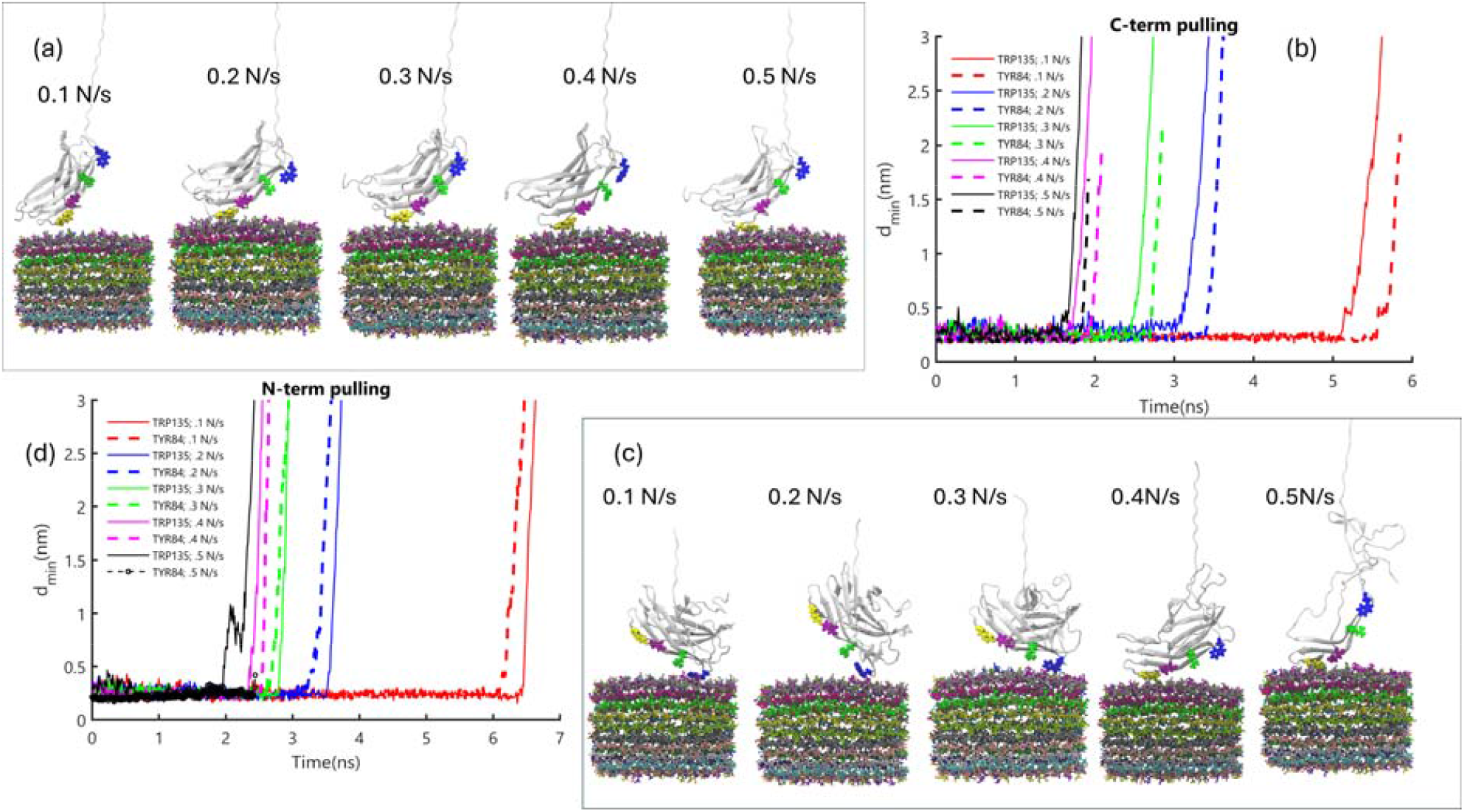
CBM3a follows a two-mode mechanism driving its unfolding/unbinding from cellulose surface: (a) CBM3a unbinding with C-terminal pulling at different loading rates. Colors: protein-white, TRP135-blue, ARG129-green, HIS74-purple, TYR84-yellow and cellulose chains in various colors, (b) The minimum distance between residues TRP135 (solid lines) and TYR 84 (dashed lines) with cellulose atoms with C-terminal pulling at different loading rates, (c) Unbinding with N-terminal pulling, (d) TRP135 and TYR84 minimum distances with cellulose atoms with N-terminal pulling.

### 3.2 Protein Structure Network Analysis

The unfolding of CBM3a when pulled by either end is governed by the protein network structure, i.e., the nature of the strongly inter-connected residue ‘communities’ within the globular structure. **Figure 5** shows the results of the network analysis of the CBM3a protein, performed over the final 5 ns of the 10 ns equilibration trajectory. Communities that consist of more than 1% of the total protein residues (2 residues) are considered. A total of 7 communities, indexed from 0 to 6, comprised of a total of 162 residues are identified and are depicted in different colors in **Figure 5a** (note that these indices are not in the order along the sequence). The remaining 8 residues at the C-terminal are found to have no community structure and are colored white. The larger communities (indices 0,1, and 3) are located closer to the N-terminus and the cellulose surface **(Figure 5a-b)** and exhibit a high number of intra-community contacts per residue 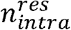 or high edge densities within the community **(Figure 5c)**. One small community at the C-terminal end (index 2) has a high number of intra-residue contacts, whereas another community in this region (index 6) has a larger number of residues but fewer intra-residue contacts. The number of inter-community contacts, as shown in **Figure 5d**, indicates strong interconnections between communities 0,1, 3, and 6. Overall, the network analysis predicts strong interconnections between the CBM3a residues located near the celF surface and the N-terminus.

**Figure 5.**
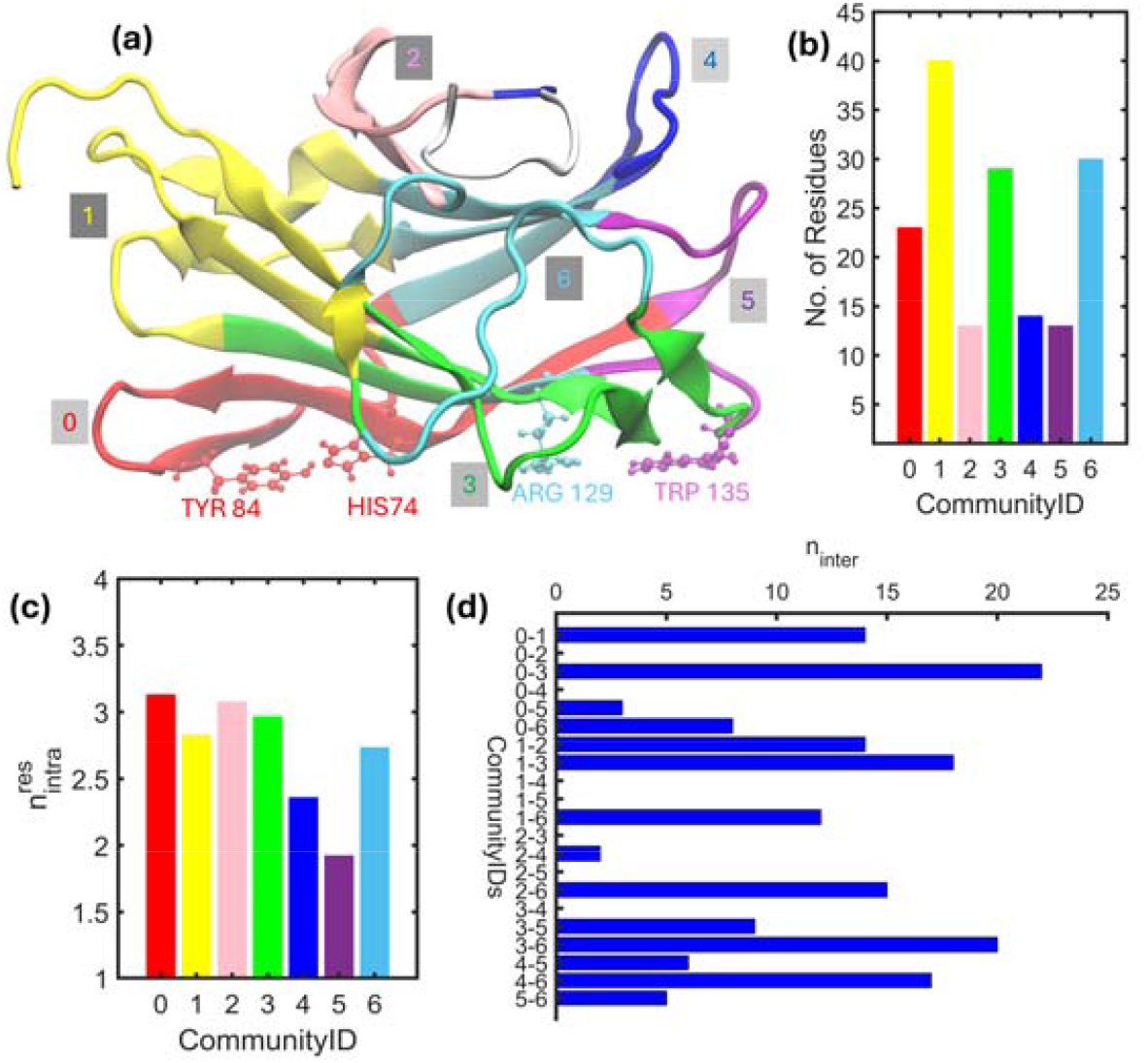
Strongly Connected Residue Communities Seen in Equilibrated CBM3a Structure Bound to Cellulose. (a) The seven communities identified through the network analysis represented in different colors, while the flexible C-terminal end colored in white does not have a community structure. The key interfacial residues TYR84, HIS 74, ARG 129 and TRP135 are indicated in ball and stick representation colored according to the community they belong to. (b) The number of residues in each community. (c) The number of intra community contacts per residue. (d) The number of intercommunity contacts between each pair of communities.

The protein residues network structure strongly influences the unfolding and unbinding behavior of CBM3a during pulling. In the case of C-terminal pulling, the extension curves at all loading rates show a two-step behavior **(Figure 3c)**. In **Figure 6a**, this behavior is analyzed at the loading rate 0.5 N/s. The linear-like increase in extension up to point 1 (550 ps) is due to the extension of the flexible region (white), which involves no unfolding. The plateau region between points 1 and 2 (550–1000 ps) indicates that although the force is increasing, not much extension occurs due to strong links that hold the community structure, preventing unfolding. Visual inspection of the snapshot at 1000 ps reveals that the step-like increase in the extension at this point is due to the breaking of community 4 (shown in blue), which required an increase in force from about 280 pN to 510 pN (approximately 230 pN for this rupture). Note that the force required to rupture the community structure is also loading rate-dependent, as inferred from **Figure 3c**. The stepwise behavior between points 2 and 3 (1000 ps −1500 ps) is due to the complete unwinding of community 2 (pink, see snapshot at 1500 ps). Further increases in force result in the partial unwinding of communities 1(yellow), 6(green), and 3 (cyan), before completely unbinding from cellulose in the TRP mode as described earlier.

**Figure 6.**
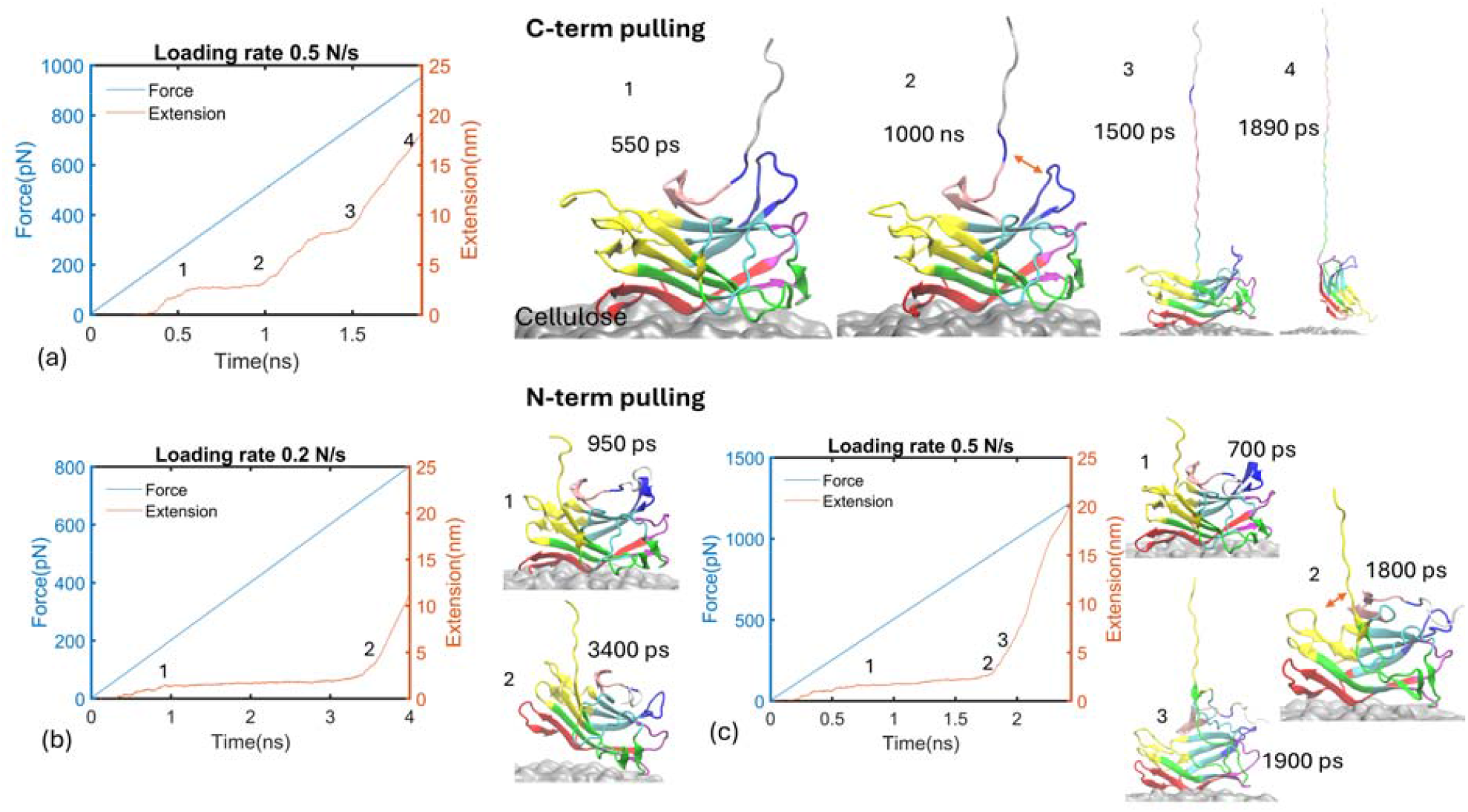
Influence of protein network structure on protein unfolding/unbinding from cellulose surface upon pulling from different ends, (a) Force/extension vs time curves during C-terminal pulling at loading rate 0.5 N/s and community structures of the protein configurations during unfolding, (b-c) Similar data in simulations with N-terminal pulling at loading rates (b) 0.2 N/s and (c) 0.5 N/s. Community color schemes are the same as in Figure 5.

**Figure 7.**
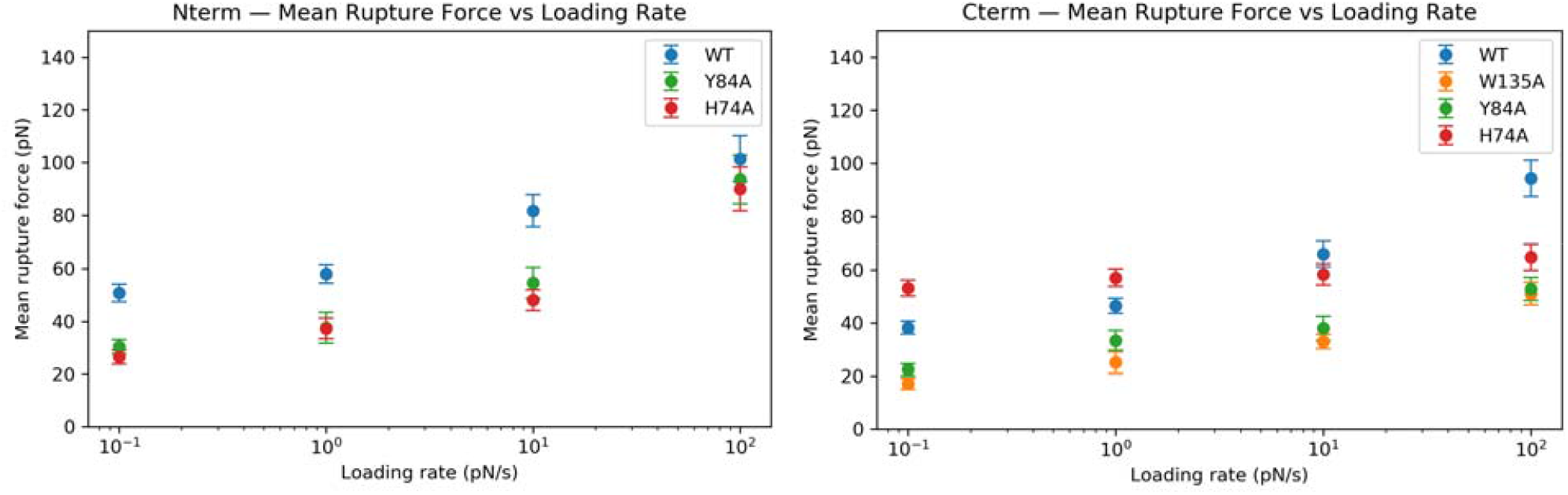
CBM3a variants pulled from its N-terminus when bound to cellulose showed higher experimentally measured rupture forces than when pulled from its C-termini. Mean rupture forces for each loading rate and CBM3a variant (WT or wild type (WT) in blue, Y84A in green, H74A in red, and W135A in orange). Here, N- and C-termini direction pulling data is shown on the left and right panels, respectively.

The N-terminal region of the protein belongs to the largest community group, community 1 (yellow). At low loading rates, this community structure is strongly held, resulting in the plateau region between points 1 and 2 in the extension curve shown in **Figure 6b**. Eventually, the protein unbinds in the TYR mode, without substantial changes in the structure of community 1. This is due to the strong interconnections between community 1 and community 0 (red), as illustrated in **Figure 6d**. These strong intercommunity contacts suggest that the TYR mode occurs even though the pulling is not directly applied to the TYR residue region. At higher loading rates **(Figure 6c)**, however, the structure of community 1 ruptures (as shown in snapshots at 1800 ps and 1900 ps), and the protein unbinds in the TRP mode, as the TYR84 region no longer feels the pull due to the rupture of the community structure. Overall, the community structure of CBM3a protein provides further insight into the unbinding mechanisms of the CBMs from cellulose surface under applied force.

### 3.3 AFS Pulling Experiments for Unbinding CBM3a from Cellulose Fibril Surface

The mechanical stability of CBM3a binding to the cellulose fibril surface was quantified using acoustic force spectroscopy (AFS). Force-induced dissociation experiments were performed on four constructs pulling from both the N and C terminus: wild-type (WT) CBM3a and three alanine-substitution variants (Y84A, H74A, and W135A). Rupture forces were measured over a range of loading rates spanning four orders of magnitude (0.1pN/s, 1pN/s, 10pN/s, and 100pN/s) for each construct. See **Supplementary Information Figure S2 and Table S1** for representative AFS rupture force datasets and statistics. Initial binding frequency differed between various constructs and loading rate conditions. See **Supplementary Information Figure S3** for representative AFS binding frequency datasets. For W135A under N-terminal pulling, initial binding was observed in approximately 15–20% of measurements (data not shown) indicating very weak binding. In contrast, all other constructs and pulling geometries exhibited initial binding frequencies between 90–100%. Bare streptavidin control beads (i.e., no GFP-CBM proteins on beads) showed low nonspecific binding, with initial binding frequencies between 0–15%.

Across all loading rates tested, the W135A variant exhibited the lowest rupture forces under both N-terminal and C-terminal pulling conditions. The Y84A and H74A variants showed intermediate rupture forces relative to WT, with smaller reductions compared to W135A. In most cases, WT CBM3a displayed the highest rupture forces at all loading rates. Rupture forces for the CBM3a variants clearly differed as a function of pulling direction. For each construct, C-terminal pulling resulted in lower rupture forces compared to N-terminal pulling at comparable loading rates. This difference was most pronounced for the WT and Y84A constructs, while the effect was smaller but still seen for H74A and W135A. Rupture force versus loading rate data were fit to the Bell–Evans model (Eq. 2) to extract apparent unbinding kinetic off-rates (k_off_) and transition state distances (Ax’). All datasets were well described by the model except for H74A under C-terminal pulling, which did not yield a reliable fit. **Table 1** summarizes the predicted unbinding off rates for all CBM3a variants when pulled from the N-versus C-terminal.

**Table 1:** Unbinding bond lifetimes predicted at zero-loading rate for CBM3a variants pulled via N-versus C-termini from cellulose. Bell Evans model fitted to experimental AFS assay rupture force data allowed determination of apparent unbinding off rate (k_off_) and transition state distance (Ax’) for each CBM3a variant.

| Bell Evans Model Predicted Apparent Bond Lifetimes |  |  |  |
| --- | --- | --- | --- |
| Terminus | Variant | $\Delta x'$ (nm) | $k_{off}$ ($s^{-1}$) |
| C-Term | WT | 0.507 | 2.15E-04 |
|  | Y84A | 0.992 | 1.18E-04 |
|  | H74A | N/A | N/A |
|  | W135A | 0.871 | 8.74E-04 |
| N-Term | WT | 0.543 | 2.87E-05 |
|  | Y84A | 0.461 | 8.56E-04 |
|  | H74A | 0.473 | 1.13E-03 |
|  | W135A | N/A | N/A |

Overall, the trends observed from experiments were consistent across the full range of loading rates tested. Briefly, rupture force measurements show a clear dependence on both CBM3a binding face aromatic residue type and pulling geometry, with W135A producing the largest decrease in rupture force and C-terminal pulling yielding consistently lower rupture forces than when pulled from the N-terminal.

## 4. Conclusions

In this study, we performed steered MD simulations and AFS experiments to study the unbinding behavior of CBM3a from cellulose surface, finding evidence that supports previously proposed multimodal unbinding modes being driven by specific CBM3a planar binding residues. Additionally, SMD simulations were performed to investigate the molecular mechanisms of CBM3a unbinding from cellulose surface, when pulled from either its C- or N-terminus of the protein to provide evidence for the very first time that pulling CBM3a from either terminus would result in very different binding properties. Our SMD results predicted that CBM3a undergoes considerable unfolding when pulled from its C-terminus, before completely detaching from the cellulose surface. In contrast, when it was pulled from the N-terminus, the protein unbinds from the surface without any unfolding. The molecular details of the unbinding process from analysis of SMD results revealed two distinct mechanisms for detachment of CBM3a from the cellulose surface: in the TRP mode, unbinding begins with the detachment of the TRP (i.e., Tryptophan 135) residue at one edge of the CBM3a planar binding motif that is engaged with the cellulose interface, whereas in the TYR mode, it initiates with the detachment of the TYR (i.e., Tyrosine 84) residue at the opposite edge of the binding motif. The SMD simulations offer stronger evidence to support predictions made based in our previous single-molecule force spectroscopy experiments and equilibrium MD simulations of CBM3a-cellulose interactions.^9^ We also observed that in the TRP mode unbinding always involves protein unwinding/unfolding, while the TYR mode unbinding occurs with protein remaining largely intact. Furthermore, based on protein network analysis of the SMD generated data, the unfolding events were seen to be clearly linked to the breaking of the internal network and community structure of the protein that further provided insights into why CBM3a unfolds when pulled from its C-termini.

The AFS results provide strong experimental evidence for the distinct unbinding modes observed in SMD simulations that are dependent on the CBM3a pulling direction. Our effector bead force spectroscopy experiments were designed to mimic cellulolytic bacteria displaying multiple CBM3a proteins tethered to the surface either via its N- or C-terminus to explore the predictions made by SMD simulations. Interestingly, our experiments revealed that the rupture force distributions and binding curves consistently showed higher mean rupture forces when CBM3a tethered to the beads was pulled off the cellulose surface from its N-terminal across all loading rates tested. This indicated a greater mechanical resistance to CBM3a when unbinding from cellulose if pulled from its N-terminal, likely due to absence of CBM3a unfolding unlike when pulled from C-terminus. We further fitted our data of rupture forces vs loading rates to the Bell Evans model to extrapolate apparent unbinding rates and transition state distances and for all mutants found that C-terminal pulling resulted in faster unbinding and lesser malleability (lower transition state distance). In the TRP-mode mediated pathway favored under C-terminal pulling, the protein is predicted by SMD simulations to undergo partial unfolding before unbinding, which is reflected experimentally for both pulling directions with lower rupture forces for the TRP to ALA mutations. In the TYR-mode mediated pathway favored under N-terminal pulling, unbinding is predicted by simulations to occur with the protein largely intact and is potentially consistent experimentally with shorter transition distances associated with lower rupture forces and higher off-rates. Experimentally, TYR and HIS mutations differed from the wild type substantially less than the TRP mutant Interestingly, TYR mutation caused a nearly 30-fold reduction in unbinding off rate when the protein is pulled from the N-terminus, with marginal difference for the same mutant when pulled from the C-terminus. Overall, there is clear agreement seen in the impact of orientation-dependent mechanical response of CBM3a as reported using various metrics in theory, experiment, and simulations. However, there is need to run additional single-molecule experiments as well as SMD simulations at comparable loading rates to experiments to further validate our findings highlighting directionality preferences for faster CBM3a unbinding as well as protein unfolding.

The fact that C-terminal pulling showed a clear impact on CBM3a unbinding as illustrated by weaker bond lifetimes (and lower mean rupture forces), potentially correlated to partial unfolding of the protein, raises the question of the biological relevance for these differences. It is likely that the CBM3a unfolding phenomenon occurring with C-terminal pulling can enable a more efficient surface diffusion of the larger cellulosomal enzyme complex to target new substrate sites and enable more efficient biomass degradation without disengaging the bacterial cell from the cellulose surface. Interestingly, for the native *aipA* scaffoldin gene, the CBM3a is expected to be tethered to the cell surface from its C-terminus and has a higher fraction of dockerin-cohesin docked enzymes that likely apply a greater force on the CBM3a on its C-terminus. By having evolved a CBM3a structure that is more malleable when pulled from its C-terminus, and easily unbind from cellulose, would give the cellulolytic bacteria a potential evolutionary advantage to more effectively grow on cellulosic biomass in a complex microbiome. Force sensitive protein engineering could be a novel approach to developing more effective enzyme/microbial biocatalysts for accelerated mechanoconversion of insoluble biomass feedstocks (e.g., terrestrial lignocellulose like corn stover or marine seaweeds like *Sargassum)*. Future studies will need to better understand the role of CBM3a unfolding and the associated cellulosomal complex activity of cellulolytic microbes like clostridia on cellulosic biomass.

## 5. Acknowledgements

SPSC acknowledges support from the National Science Foundation (NSF CBET CAREER Award 1846797).

