## Supporting Information for "Force-induced unfolding versus unbinding of a cellulosomal carbohydrate-binding module from cellulose interface"

**Additional Supporting Data Provided in SI as Separate Files**

### 1. PDB files of CBM3a–Cellulose configurations from SMD simulations (CBM3aCellulosePDBs.zip)

2. Gene sequences for all constructs tested (Pulling Direction SI Nucleotides-2-FINAL.xlsx)

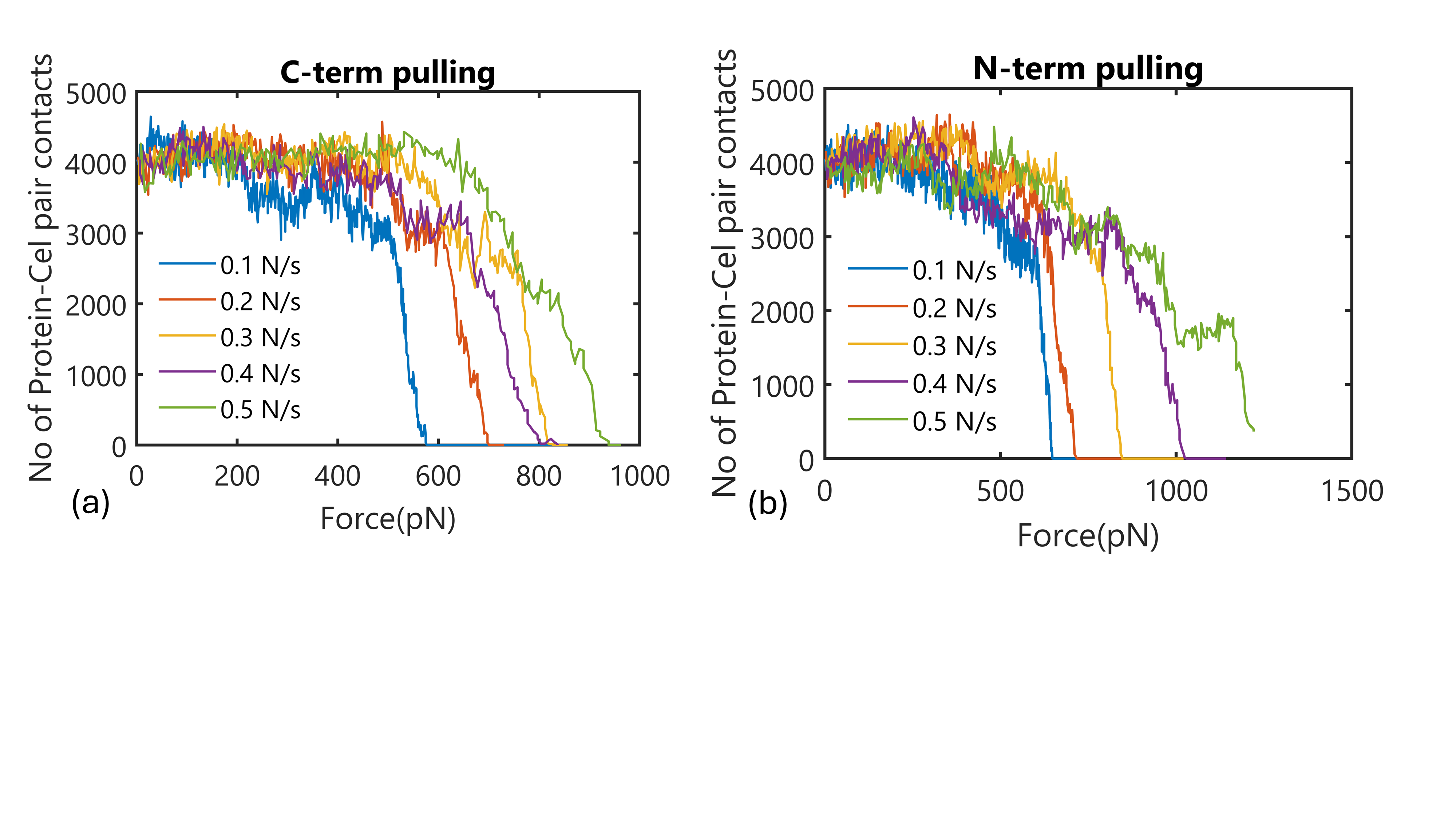

**Figure S1.** **The number of protein-CelF atomic pair-contacts in SMD simulations.** Here, (a) C-terminal pulling and (b) N-terminal pulling. The contacts are calculated with the pair -distance less than 0.6 nm.

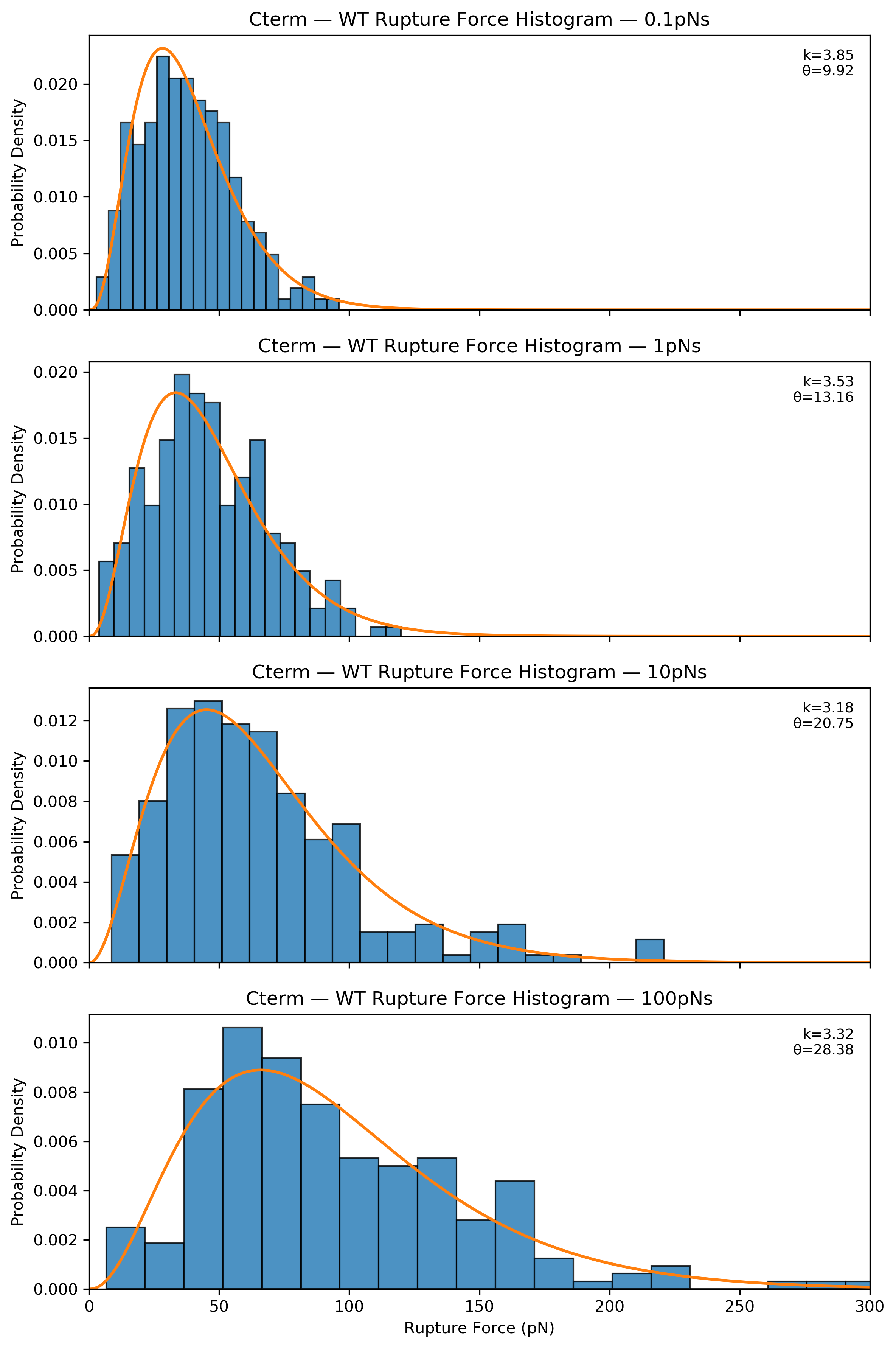

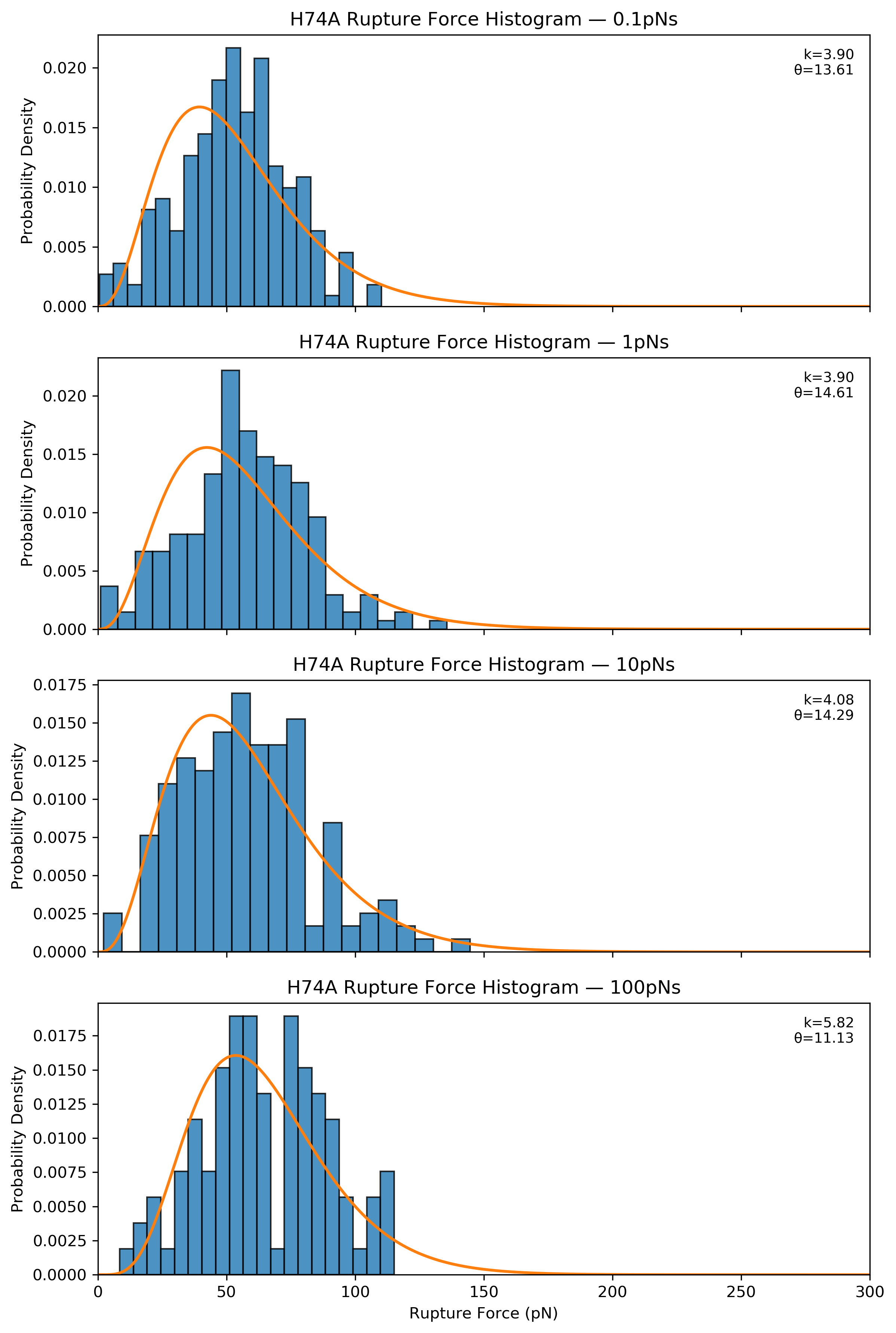

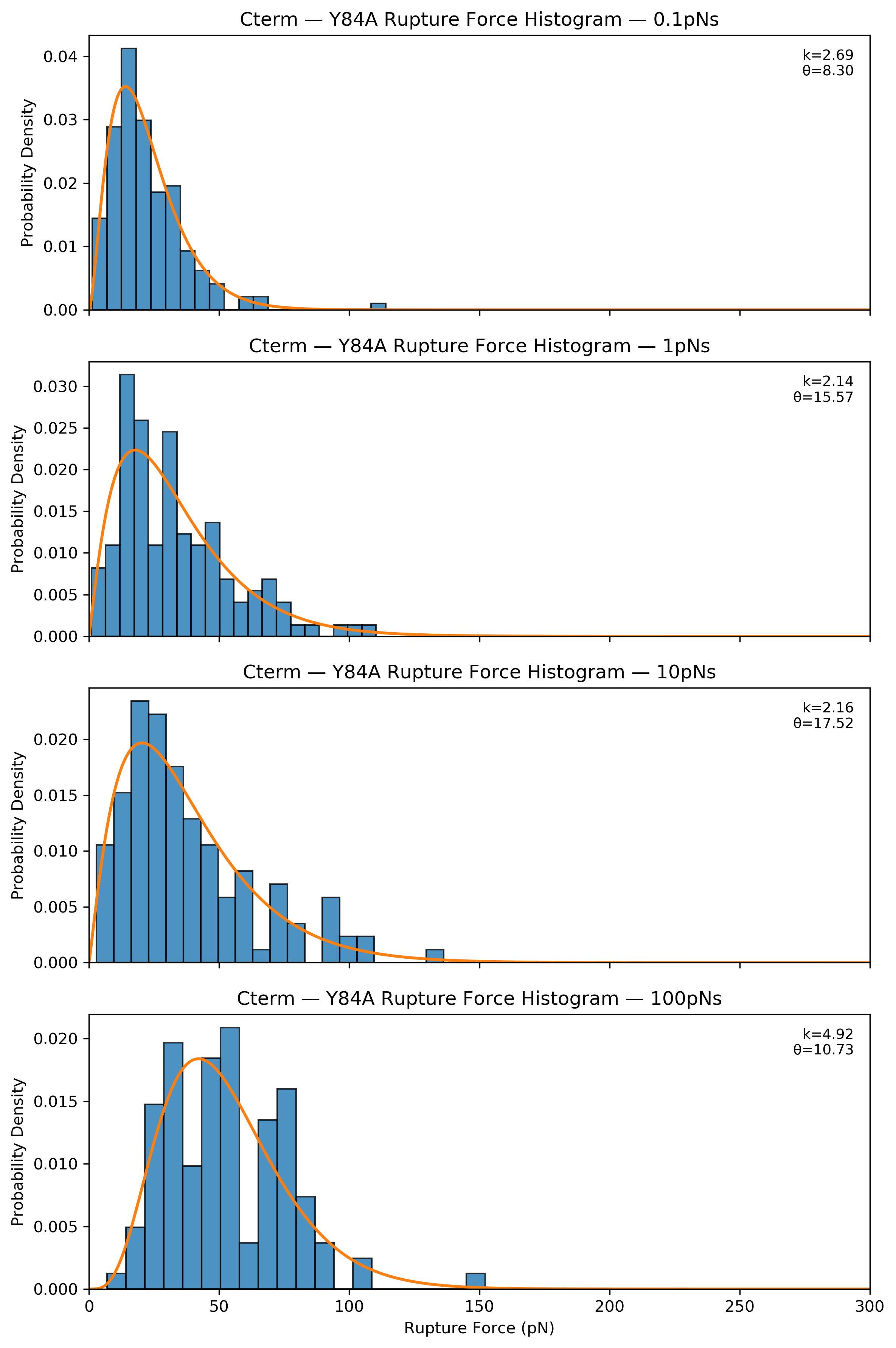

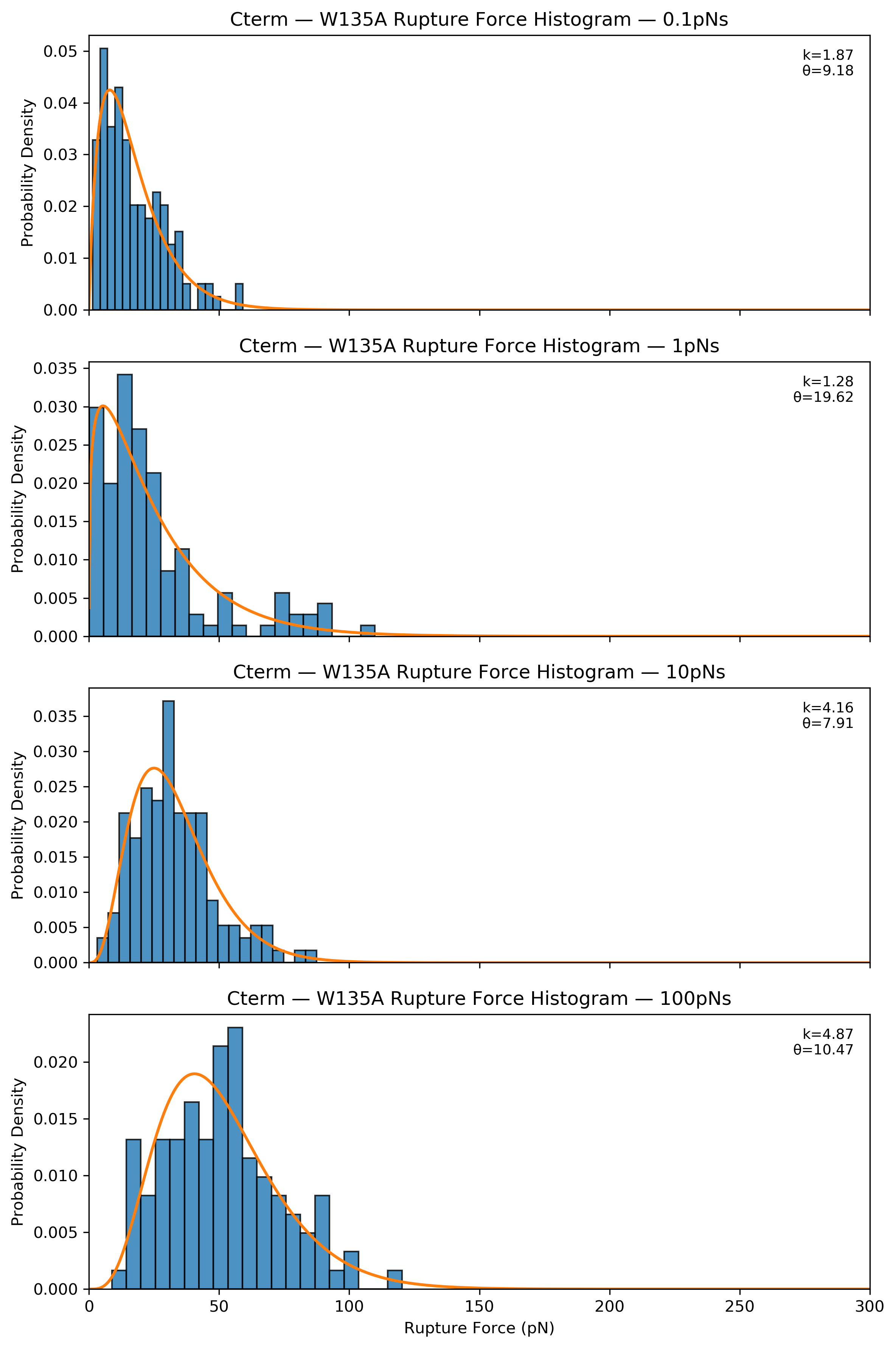

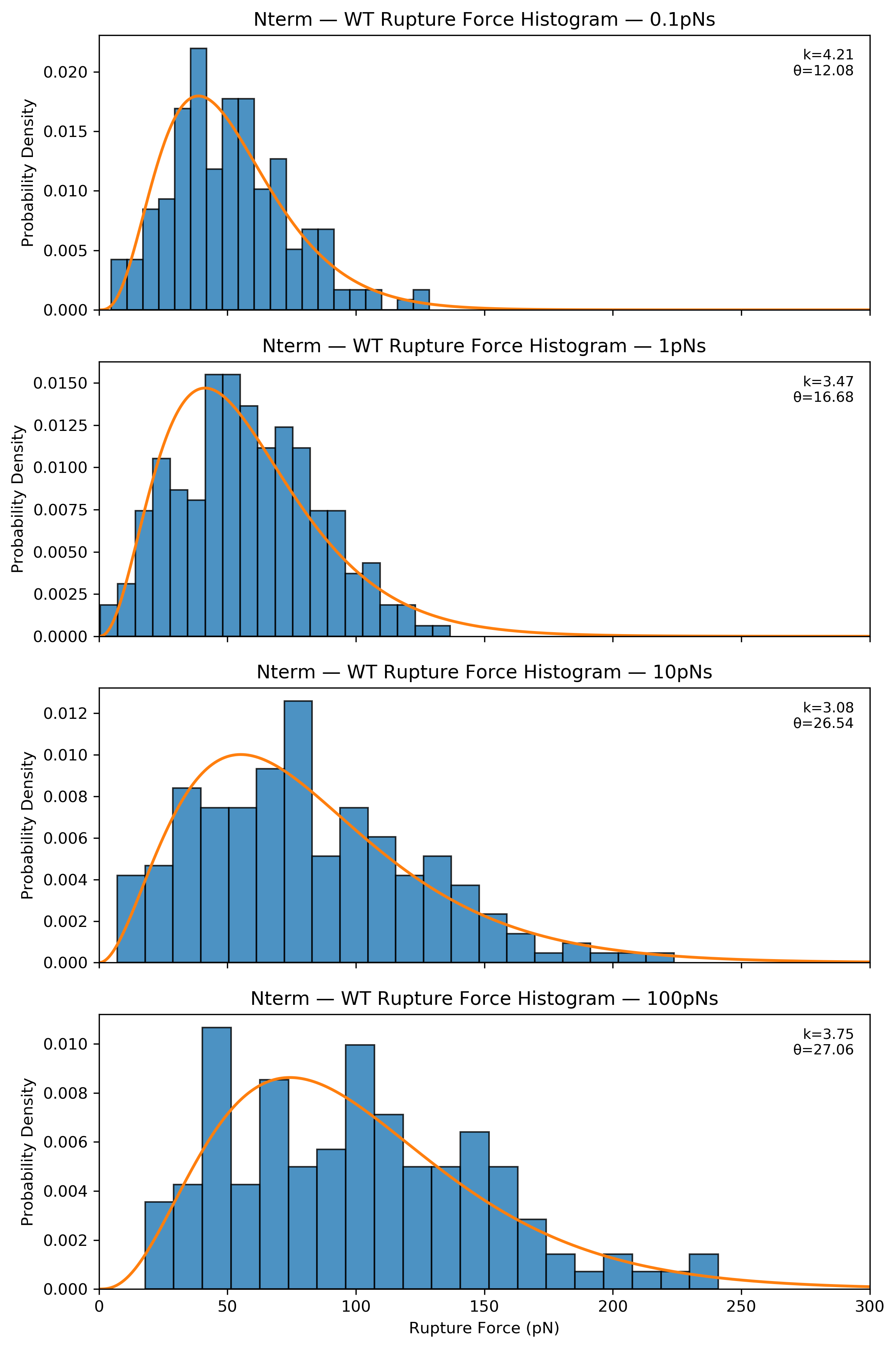

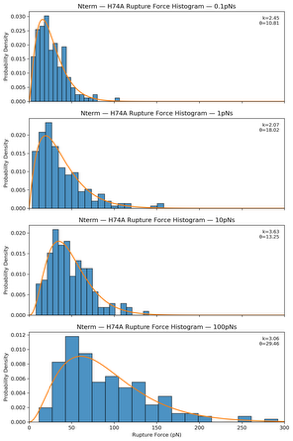

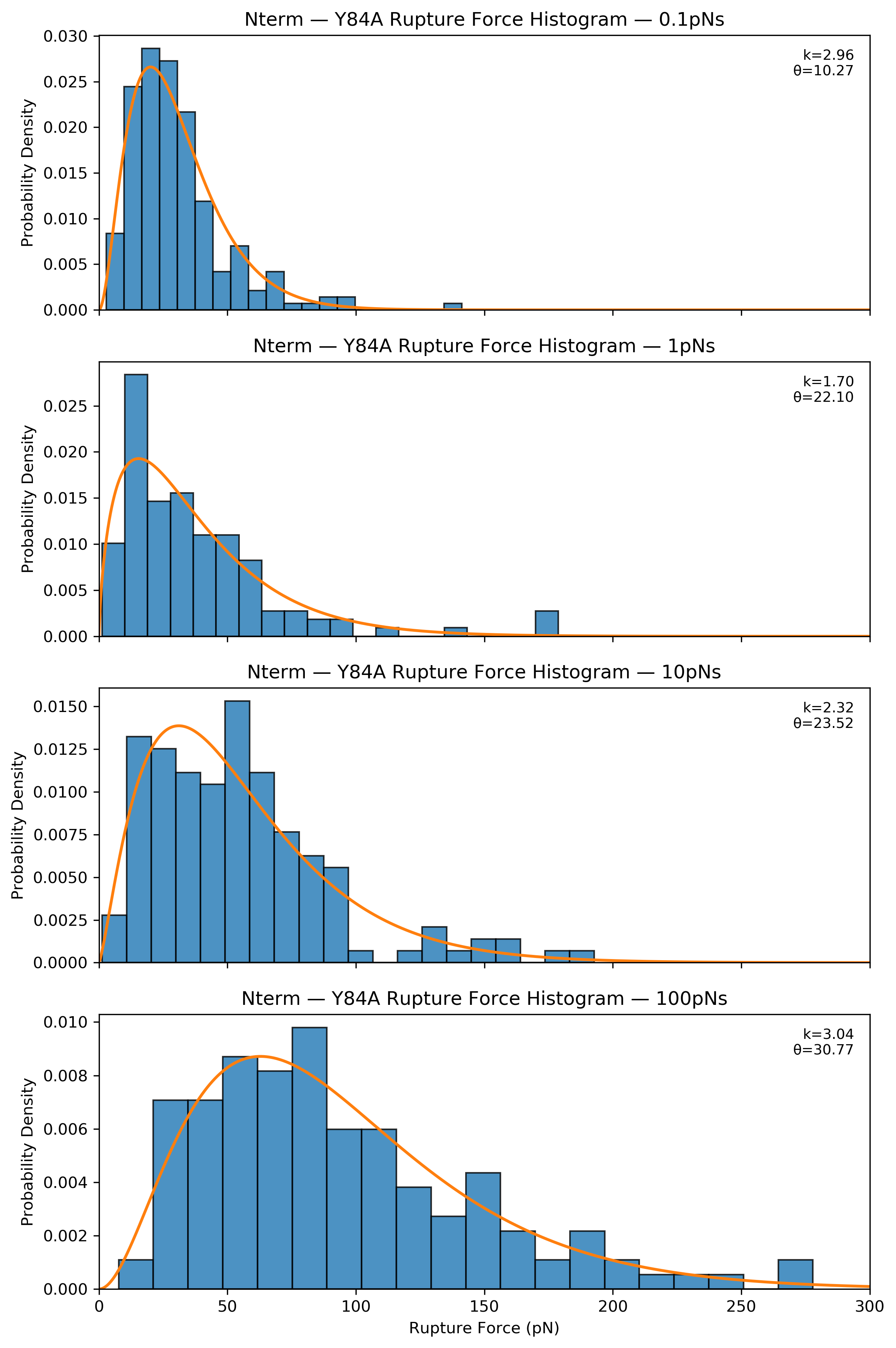

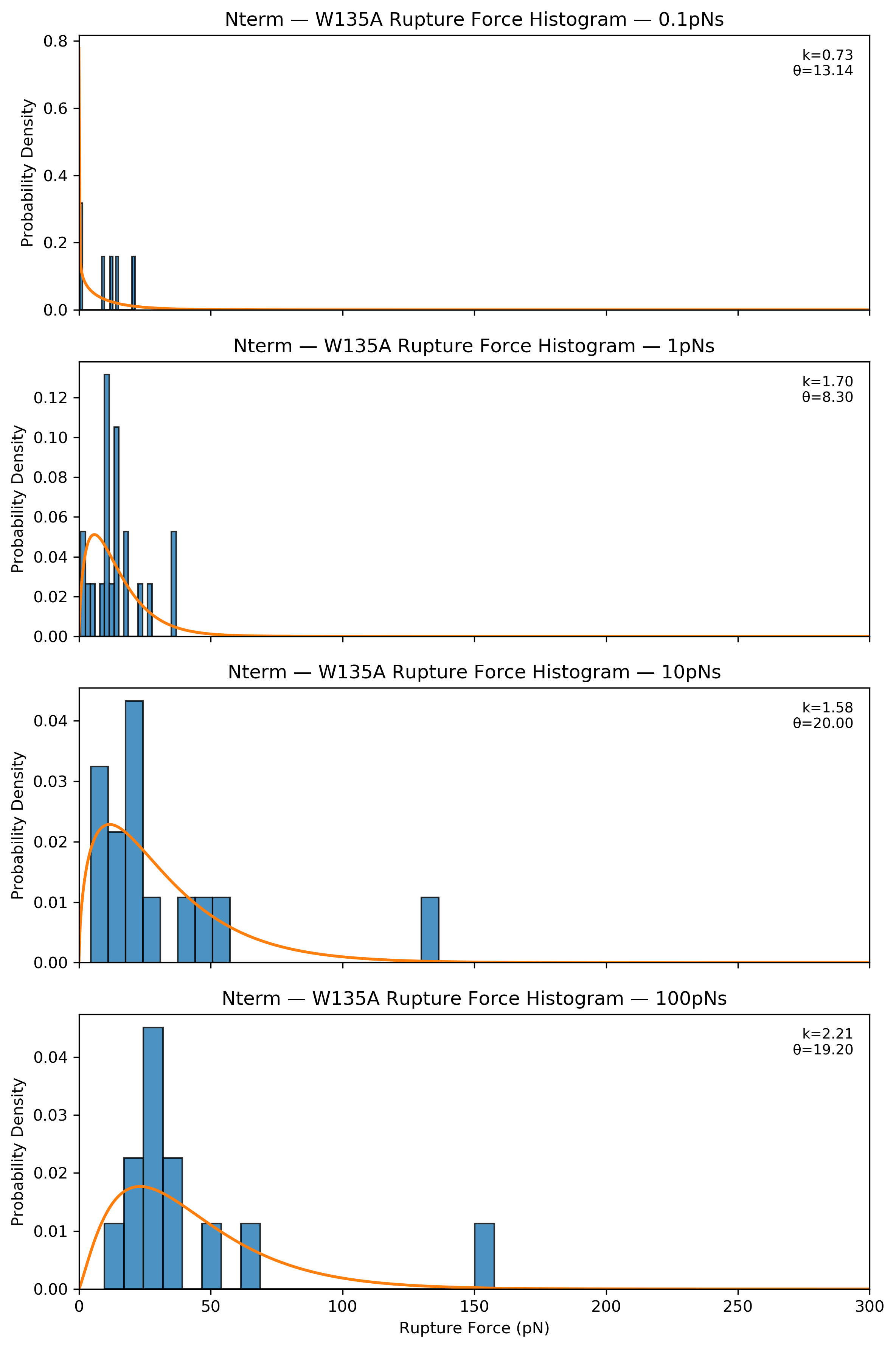

**Figure S2.** **Representative Histograms of Rupture Force Distributions from AFS CBM3a-Cellulose Pulling Assays**. Here, Top Row: C-Terminal Pulling, Bottom Row: N-Terminal Pulling. First Column: WT-CBM3a, Second Column: H74A-CBM3a, Third Column: Y84A, Fourth Column: W135A. In descending order (top to bottom) each of the subsections are loading rates of 0.1pN/s, 1pN/s, 10pN/s, and 100pN/s.

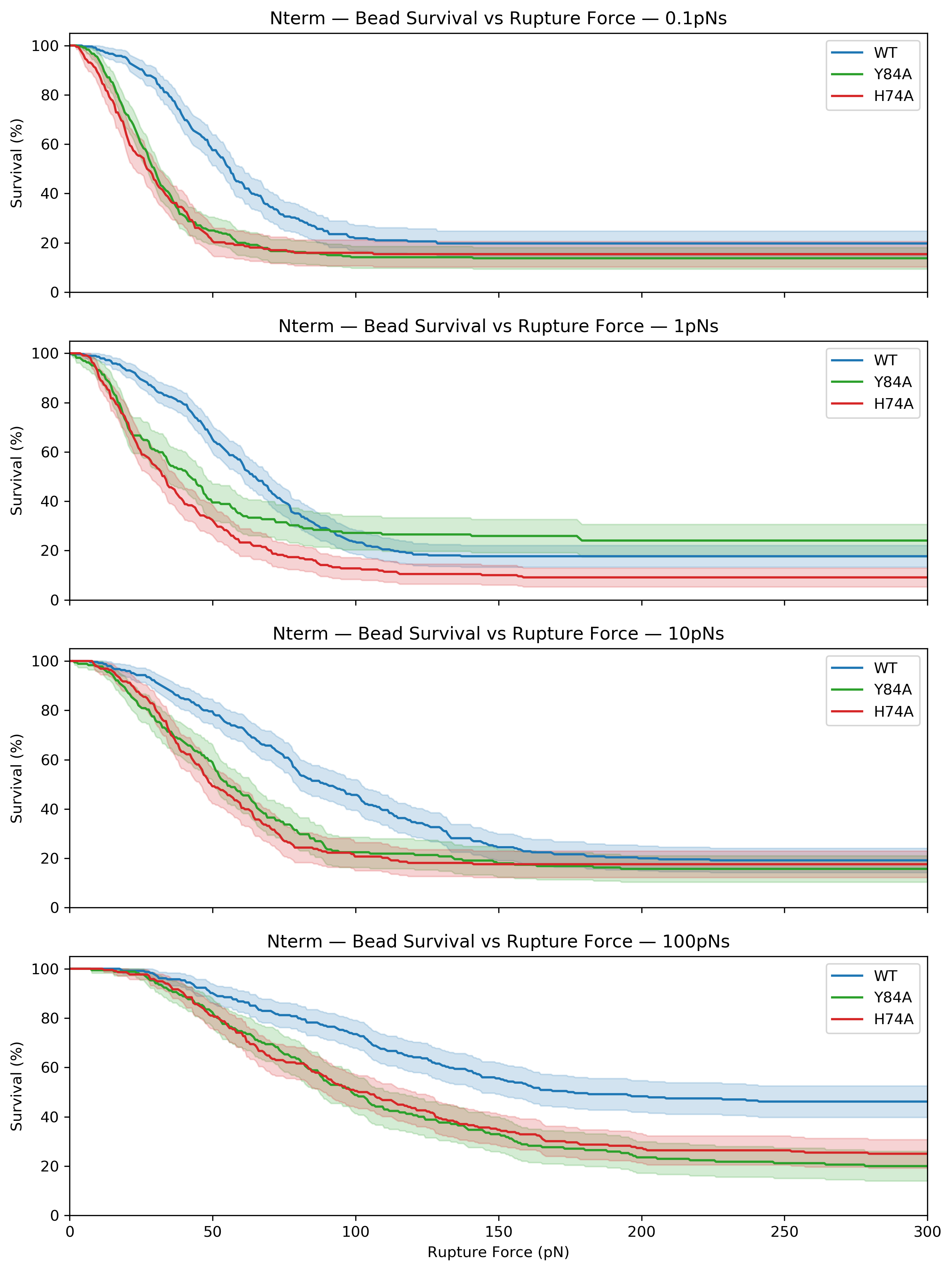

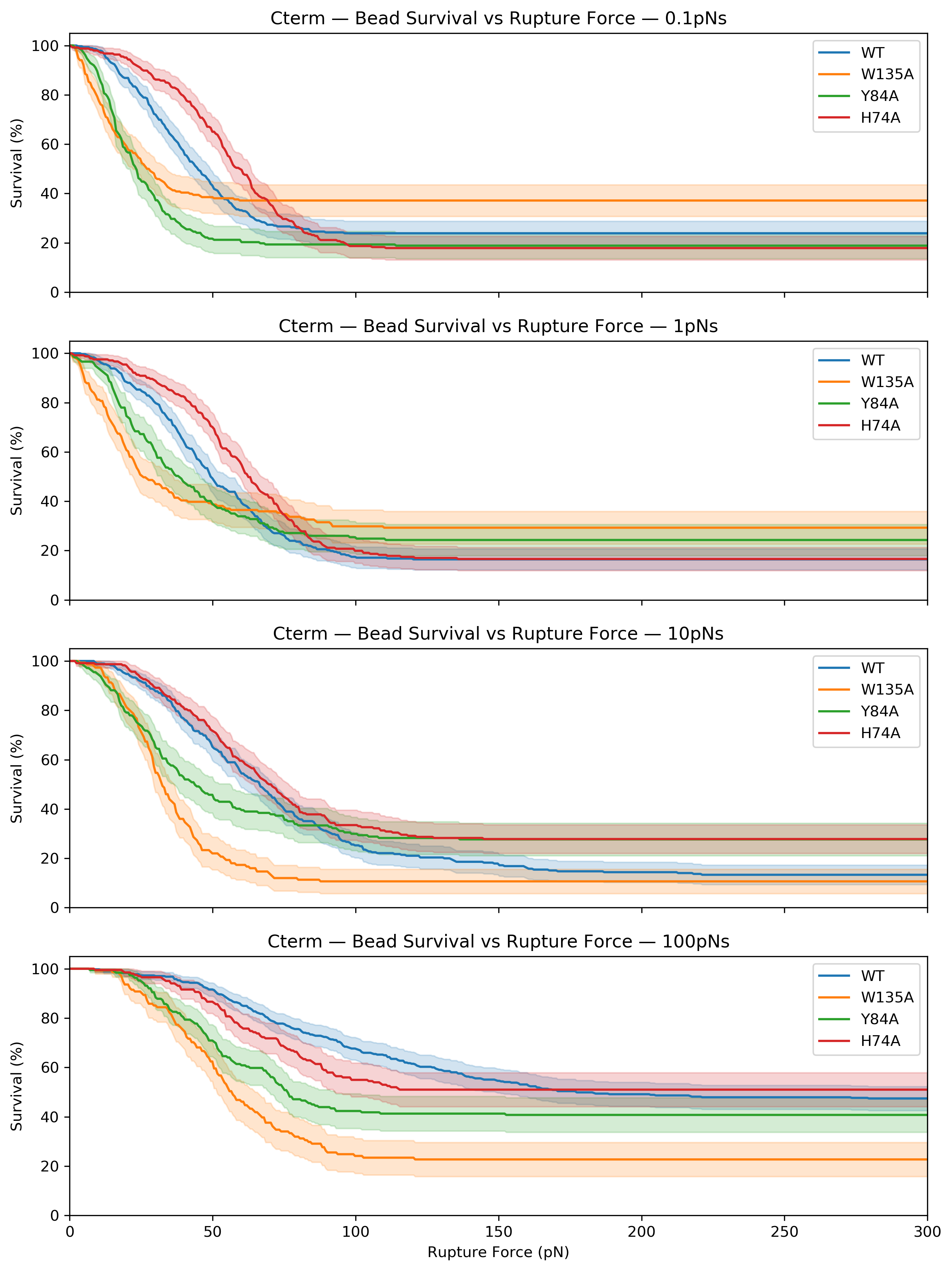

**Figure S3. Percentage of CBM3a tethered beads remaining bound to cellulose surface as function of applied force for each CBM3a variant (WT in blue, Y84A in green, H74A in red, W135A in orange)**. Here, Left column: N-terminal pulling, Right column: C-terminal pulling. In descending order *(top to bottom)* each of the subsections are loading rates of 0.1pN/s, 1pN/s, 10pN/s, and 100pN/s.

| **N Term** | | | | | | |
| --- | --- | --- | --- | --- | --- | --- |
| **Variant** | **Condition** | **N_values** | **Mean** | **StdDev** | **CI95_Lower** | **CI95_Upper** |
| H57A | 0.1pNs | 159 | 26.48782 | 16.85805 | 23.84726 | 29.12838 |
| H57A | 1pNs | 199 | 37.31826 | 28.20574 | 33.37531 | 41.26122 |
| H57A | 10pNs | 159 | 48.05432 | 24.87457 | 44.15809 | 51.95055 |
| H57A | 100pNs | 163 | 90.08872 | 53.99228 | 81.73764 | 98.43979 |
| Y67A | 0.1pNs | 207 | 30.39057 | 19.30188 | 27.7456 | 33.03554 |
| Y67A | 1pNs | 123 | 37.55275 | 32.70078 | 31.71584 | 43.38966 |
| Y67A | 10pNs | 150 | 54.59388 | 35.93698 | 48.79578 | 60.39198 |
| Y67A | 100pNs | 136 | 93.60606 | 54.21057 | 84.41272 | 102.7994 |
| W118A | 0.1pNs | 6 | 9.607267 | 8.177377 | 1.025636 | 18.1889 |
| W118A | 1pNs | 21 | 14.1478 | 9.797405 | 9.688078 | 18.60753 |
| W118A | 10pNs | 14 | 31.6445 | 33.2916 | 12.4225 | 50.8665 |
| W118A | 100pNs | 12 | 42.3814 | 38.90484 | 17.66244 | 67.10035 |
| WT | 0.1pNs | 191 | 50.8267 | 23.46282 | 47.47792 | 54.17548 |
| WT | 1pNs | 237 | 57.97749 | 27.07895 | 54.51221 | 61.44277 |
| WT | 10pNs | 198 | 81.72371 | 43.50701 | 75.62622 | 87.8212 |
| WT | 100pNs | 126 | 101.398 | 49.85299 | 92.6082 | 110.1878 |
| **C-Term** | | | | | | |
| Variant | Condition | N_values | Mean | StdDev | CI95_Lower | CI95_Upper |
| H57A | 0.1pNs | 202 | 53.12227 | 21.38741 | 50.15502 | 56.08952 |
| H57A | 1pNs | 201 | 56.99025 | 23.68099 | 53.69653 | 60.28396 |
| H57A | 10pNs | 166 | 58.24697 | 26.0718 | 54.25155 | 62.24238 |
| H57A | 100pNs | 99 | 64.73323 | 24.46274 | 59.85422 | 69.61224 |
| Y67A | 0.1pNs | 172 | 22.30855 | 14.47557 | 20.12981 | 24.48728 |
| Y67A | 1pNs | 134 | 33.31352 | 21.6764 | 29.60967 | 37.01736 |
| Y67A | 10pNs | 128 | 37.90331 | 26.25585 | 33.31105 | 42.49558 |
| Y67A | 100pNs | 112 | 52.7994 | 23.43364 | 48.41167 | 57.18713 |
| W118A | 0.1pNs | 137 | 17.16676 | 12.27096 | 15.09352 | 19.23999 |
| W118A | 1pNs | 128 | 25.02309 | 23.37309 | 20.93502 | 29.11115 |
| W118A | 10pNs | 134 | 32.90453 | 15.84057 | 30.19786 | 35.61121 |
| W118A | 100pNs | 109 | 51.00126 | 22.10059 | 46.80529 | 55.19723 |
| WT | 0.1pNs | 220 | 38.14395 | 18.21726 | 35.72333 | 40.56457 |
| WT | 1pNs | 244 | 46.49214 | 22.65735 | 43.635 | 49.34927 |
| WT | 10pNs | 247 | 65.91119 | 38.48083 | 61.08854 | 70.73385 |
| WT | 100pNs | 214 | 94.34907 | 50.74351 | 87.51159 | 101.1866 |

**Table S1**: **Summary Table of Mean Rupture Forces** **from AFS data Reported in Fig S2**. Here, Standard Deviations (StdDev), Confidence intervals (CI95), and Number of data points (N_values), for N- (top) and C- (bottom) terminal pulling data are reported.
