## Supporting Information for "Force-induced unfolding versus unbinding of a cellulosomal carbohydrate-binding module from cellulose interface"

### PDB Files of CBM3a–Cellulose Configurations from the Simulations

The PDB files of the initial and final configurations of the CBM3a–cellulose system obtained from the simulations at different loading rates and pulling modes are provided.

The initial configuration for all simulations is a structure of CBM3a adsorbed on a cellulose fiber after 10 ns of equilibration, provided as **CBM3aCellulose.pdb**.

The final configuration files are named according to the pulling mode, loading rate, and simulation time. The prefixes **CT** and **NT** denote **C-terminal** and **N-terminal** pulling, respectively, while **g** indicates the loading rate in N/s. The simulation time corresponding to the final configuration is included in the filename.

For example, **CBM3aCelluloseCTg0.5_1.92ns.pdb** corresponds to the final configuration from a simulation with **C-terminal pulling** at a **loading rate of 0.5 N/s**, where **1.92 ns** is the simulation time of the final structure.
